# Modeling Fibrosis with MASH Patient Liver-Derived Organoids

**DOI:** 10.1101/2025.09.18.677209

**Authors:** Andy Liu, A. Dylan T. Haseman, Khushal Bantu, Jia Nong, Vladimir Muzykantov, Jilian Melamed, Michael Kegel, Jenna Muscat-Rivera, Drew Weissman, David Smith, Mei Zhang, Daniel J. Rader, Tobias D. Raabe

## Abstract

Metabolic dysfunction-associated steatohepatitis (MASH) can lead to liver fibrosis and cirrhosis ultimately leading to liver transplantation or death. Therapeutic options for MASH-associated fibrosis are limited in part because of the lack of good model systems.

To address this challenge, we developed a 3D MASH liver fibrosis model by using organoids derived from MASH patient liver co-cultured with human liver-derived hepatic stellate cells (HSC) and human peripheral blood monocytes (MC). Spontaneous self-assembly resulted in fibrotic scar-like 3D structures with senescent parenchymal cells, proliferating collagen secreting myofibroblasts (MFB) and proinflammatory TREM2+ scar-associated macrophages (MP). Single cell RNA sequencing suggested high similarity with MASH patient liver fibrotic scars.

Lipid nanoparticles (LNPs) formulated with anti-*YAP1*siRNA could specifically and efficiently knockdown YAP1 in the MFBs, resulting in MFB senescence, a desirable therapeutic goal. This MASH patient liver-derived fibrosis model opens novel avenues towards testing treatments for MASH-associated liver fibrosis with reduced adverse effects.

## Introduction

Metabolic dysfunction-associated steatohepatitis (MASH), the progressive form of metabolic dysfunction-associated steatotic liver disease (MASLD), has become a leading cause of chronic liver disease worldwide. Characterized by hepatic steatosis, inflammation, and fibrosis, MASH significantly increases the risk of cirrhosis, hepatocellular carcinoma, and liver failure. Its prevalence has grown alongside metabolic syndrome, obesity, and type 2 diabetes, which affects approximately one third of the adult U.S. population. As a result, MASH cases in the United States are forecasted to increase by +82.6%, from 11.61 million (2020) to 19.53 million (2039) (1). Despite its increasing prevalence, effective pharmacological treatments remain lacking; the few FDA-approved drugs available offer modest benefits, especially in their limited ability to reduce liver fibrosis (2).

MASH is the result of a complex interplay of metabolic insults including insulin resistance, dyslipidemia, and adipose tissue inflammation leading to hepatic fat accumulation (hepatic steatosis) which is termed metabolic dysfunction-associated fatty liver disease (MAFLD). In a subset of MAFLD patients, MASH develops due to a combination of genetic and environmental factors that are not yet fully understood. Hepatocyte lipotoxicity leads to hepatocellular stress, oxidative injury, and activation of pro-inflammatory and pro-fibrotic pathways, involving Kupffer cell activation, recruitment of monocyte (MC)-derived TREM2+ macrophages (MPs), and trans-differentiation of hepatic stellate cells (HSCs) to proliferating myofibroblasts (MFBs) which produce excessive extracellular matrix, ultimately resulting in irreversible end stage fibrosis. A major limitation for the field is the lack of a valid *in vitro* model system for the transition process of MASH to fibrosis.

Here we report the use of end stage MASH patient liver-derived organoids we described previously (3) to study major hallmarks of human end stage MASH that have proven challenging to investigate with MASH animal models: 1) the long incubation time of 10-20 years, 2) the accumulation of ballooning, inflamed hepatocytes, 3) the emergence of senescent biphenotypic hepatocyte /cholangiocyte intermediates that overlap with the fibrotic strands of MFBs surrounding the regenerative nodules, and 4) the senescent cell-derived proinflammatory factors that help cause MFB proliferation and fibrosis. We took advantage of the ability of MASH patient liver-derived organoids to retain the epigenetic memory of the end stage MASH liver. In contrast, iPSC-derived hepatic organoids do not retain the epigenetic memory of the MASH liver because iPSC generation depends on complete reprogramming of blood or skin cells to the native state (4).

We showed previously that MASH liver tissues can give rise to organoids that grow and can be passaged by mechanical disassociation. The organoid cells are bipotent, and upon exposure to a defined differentiation medium, they stop dividing and then differentiate into both hepatocyte-like cells and cholangiocyte-like cells *in vitro* (3, 5). However, these organoids do not contain HSC-derived MFBs and peripheral blood-derived TREM2+ MPs, which are other key cell types that strongly accumulate in the fibrotic scar of end stage MASH liver (6, 7). To address this, we added human liver-derived HSCs and human peripheral blood-derived MCs to the MASH liver-derived organoids. Interestingly, these three cell types interact in culture to spontaneously form a MASH liver-like fibrotic scar which - by both microscopy and single cell analysis - is surprisingly similar to the fibrotic scars found in end stage MASH patient liver.

In addition, we report a novel LNP-based approach to gene silencing in MASH liver organoids with effective reduction of fibrosis. Genome editing has become an indispensable tool in disease modeling, but its application to 3D organoid systems has been hindered by technical challenges. Existing approaches typically require genome editing at the iPSC or early passage adult stem cell stage, followed by differentiation into mature organoids - an inefficient and time-consuming process. Recent advances in lipid nanoparticle (LNP)-mediated gene silencing, including successful knockdown of liver-produced transthyretin (TTR) to alleviate amyloidosis (8), use of mRNA LNPs for overexpression of immunogenic proteins, most notably for COVID19 (9), or LNP-mediated editing of the *CPS1* gene to treat severe CPS1 deficiency(10), suggest a much more efficient approach.

## Results

### Development of a human MASH liver fibrosis organoid model

To generate a MASH fibrosis model (see **Figure 1** and methods for details), we first considered which cell types to use to best compromise between representing the most important cell types involved in fibrosis and the technical demands of handling multiple cell types in the same 3D organoid structure. Based on available literature including single cell data from the human fibrotic liver (7) and our previous published work (3), we decided to use liver parenchymal cells from MASH patient liver-derived organoids, commercially available primary human liver-derived HSCs/MFBs, and primary human peripheral blood-derived MCs.

**Figure 1:**
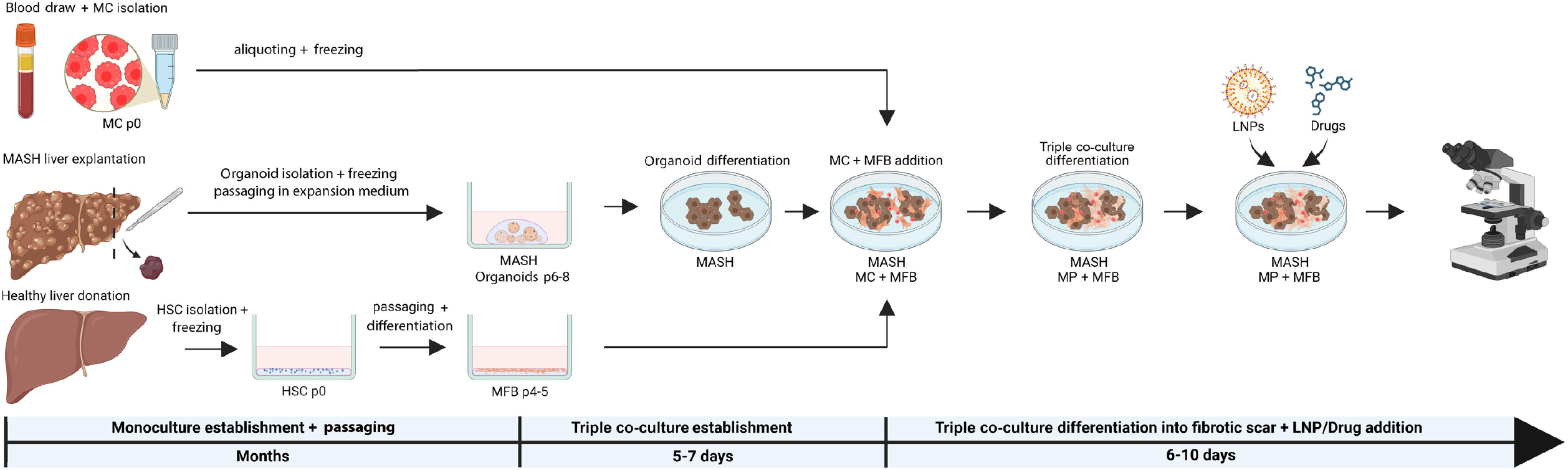
Timeline for a novel MASH patient liver-derived 3D fibrosis model. The model is assembled entirely from end stage MASH patient liver-derived hepatic organoids (MASH), healthy donor liver-derived hepatic stellate cells (HSC), and healthy human peripheral blood monocytes (MC). In our fibrosis medium, they differentiate into MASH hepatocytes, proliferative myofibroblasts (MFB), and activated macrophages (MP), respectively. After assembly the 30 fibrosis model is stable for at least 10 days.

We previously described in detail the derivation and functional characterization of organoids derived from 3 healthy donor livers and 6 MASH patient explant livers. We found that MASH organoids exhibit a senescence phenotype as they had much lower capacity to replicate than healthy organoids (3) (**Figure S1**). More recently, using the corresponding organoid RNA seq data we found that all 6 lines of our MASH organoids spontaneously over-expressed multiple senescence markers compared to the average of the 3 healthy organoid lines (**Figure S2**). Importantly, all the over-expressed senescence markers are part of the functionally annotated Sen Mayo senescence gene set (11). For this study, we chose MASH organoids from a 53-year-old male (patient 3, PAT3), selected amongst the 6 different MASH patients as they seemed to represent an ‘average’ phenotype of MASH liver organoid lines both in terms of their senescence phenotype and functional features (**Figures S1, S2**). As a control, we used organoids-derived from the liver of a 28-year-old healthy male donor (MLD), selected amongst 3 different donors as they represented the average phenotype of healthy liver organoid lines. **(Figures S1, S2)**. We obtained the HSCs isolated from human healthy donor liver at passage 1 from Sciencell, Inc and fresh healthy human peripheral MCs from the Penn Immunology Core.

We experimented with different media, plate coatings, and different ratios of the above three cell types within triple cultures, as well as different timing schedules for combining these components. The main goal was to allow for full differentiation of hepatocyte-like cells and cholangiocyte-like cells derived from the organoids while simultaneously fully differentiating peripheral MCs to TREM2*+* MPs that can physically interact with the terminally differentiated MFBs with high collagen production.

Our resulting best protocol (see **Figure 1** and methods for details) started with generating 2D layers of MASH liver-derived organoids cells to which we subsequently added human liver-derived MFBs and healthy primary human peripheral MCs. Fibrotic scar-like structures developed during the first 5 days of co-culture, and the system itself was stable for at least another 10 days afterwards. Importantly, during the first 5 days of co-culture, the MCs spontaneously differentiated into TREM2+ activated MPs that predominantly bind to fully differentiated MFBs. **Figure 1** shows the basic experimental setup and timeline for the MASH fibrosis model.

### MASH liver-derived organoid cells show fibrosis-associated phenotypes

Important properties of fully differentiated non-proliferating MASH liver organoid cells, either in monoculture or co-culture with MFBs, are shown in **Figure 2**. These cells are biphenotypic, as they express both a classic cholangiocyte marker (CK19; *KRT19*) (**Figure 2A**) and a classic hepatocyte marker (CK18; *KRT18*) (**Figure 2B**). Importantly, MASH liver-derived organoids in co-culture with MFBs form SMA-1 positive fibrotic scar-like structures (**Figure 2A, Figure 3B, D**). High magnification images show direct contact between organoid cells and MFBs forming dense fibrotic bands similar to what is seen in the natural human MASH liver. Surprisingly, MFBs growing on top of healthy liver-derived organoids show weaker SMA-1 staining and less spindle formation (**Figure S3**). Furthermore, MASH liver organoids form cells that are larger than healthy liver organoid cells in both cellular diameter and nuclear diameter, reminiscent of the ballooning of biphenotypic hepatocyte/cholangiocyte cells in end stage human MASH liver (**Figure 2B**). Finally, MASH liver-derived organoids acquire a spindle-like shape and form large stress vacuoles in response to exposure to the free fatty acids oleic and palmitic acid at concentrations present in human MASH liver (**Figure 2C**), indicating cellular stress. This phenotype occurs in MASH patient liver specifically within the fibrotic bands but not in the non-fibrotic areas **(Figure 4**). Interestingly, the healthy liver-derived organoid cells did not demonstrate this phenotype, indicating a fundamentally different physiology (**Figure 2C**). Overall, these data show that MASH liver-derived organoids produce fully differentiated cells that are similar to the ballooning hepatocytes and biphenotypic hepatocytes/cholangiocytes in human end stage MASH liver.

**Figure 2:**
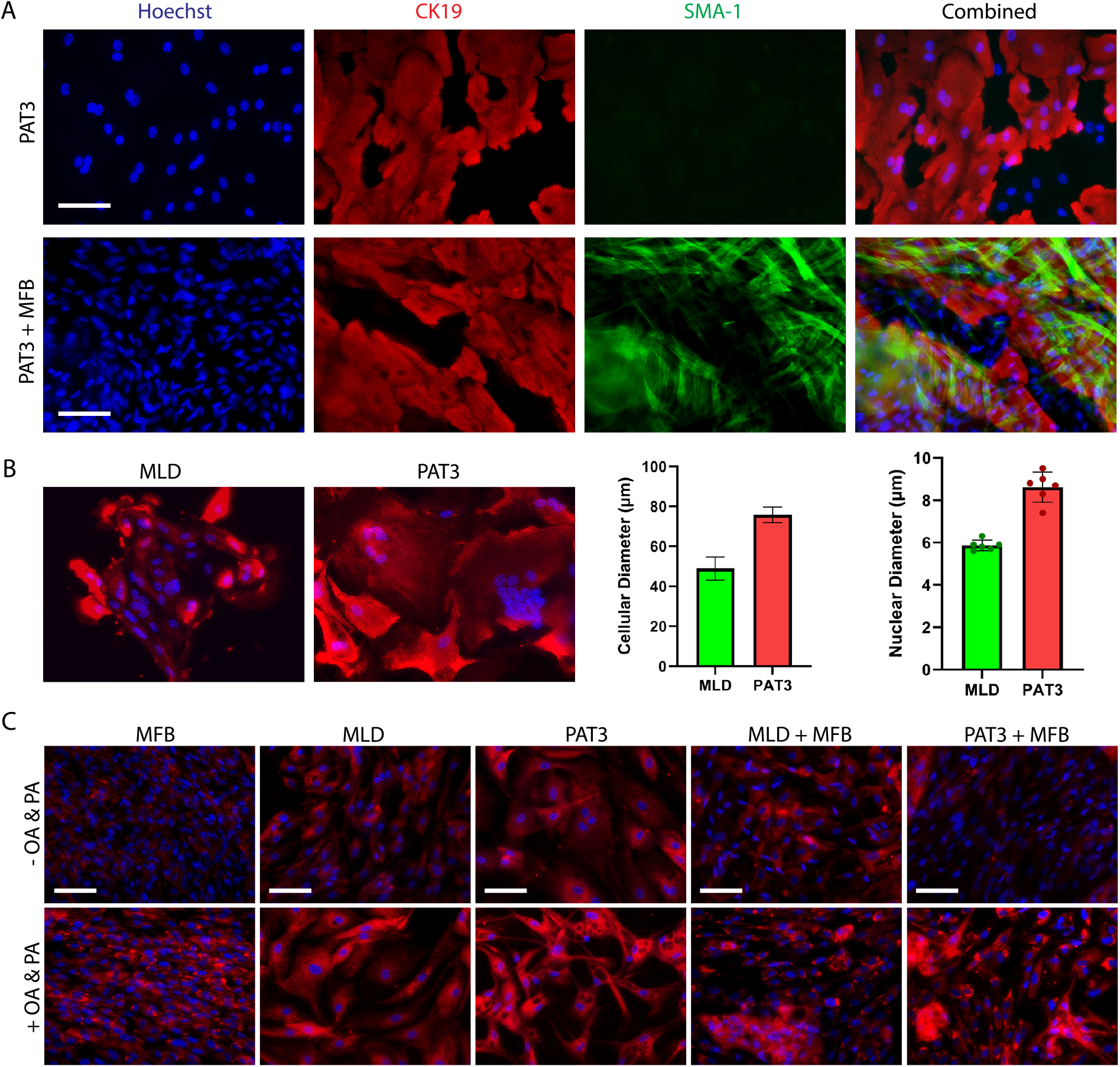
The MASH fibrosis model recapitulates important characteristics of the biphenotypic hepatocyte/cholangiocyte cells of the human MASH fibrosis. A) MASH liver-derived organoids (PAT3) express cytokeratin-19 (CK19), and MFB express a-smooth muscle actin (SMA-1). Upper panel: A PAT3 organoid monoculture is shown as a control. Lower panel: PAT3 double co-cultures with MFBs in the context of the fibrosis model. Note the strongly striated MFBs on top of the organoids. Scale bars, 100 µm. B) PAT3 double co-cultures with MFBs recapitulate the ballooning phenotype of biphenotypic hepatocyte/cholangiocyte cells of human MASH liver. PAT3s exhibit both larger cellular and nuclear diameter compared to healthy liver-derived organoids (MLD). Cells are stained with CK18. Mean ± SD, n = 91 for cellular diameter, n = 8 for nuclear diameter. C) In the presence of the free fatty acids (FFA) oleic acid and palmitic acid, triglyceride droplets form as in the natural fibrosis of the human MASH liver. PAT3s, but not MLDs, develop large vacuoles and branched limbs, indicating major cellular stress reminiscent of biphenotypic cells in fibrotic areas of the human MASH liver. Both PAT3 and MLDs accumulate triglyceride droplets in response to the FFA addition. MFBs also take up FFA across monocultures and co-cultures and form triglyceride lipid droplets. Triglyceride lipid droplets are visualized by LipidTox staining (red).

**Figure 3:**
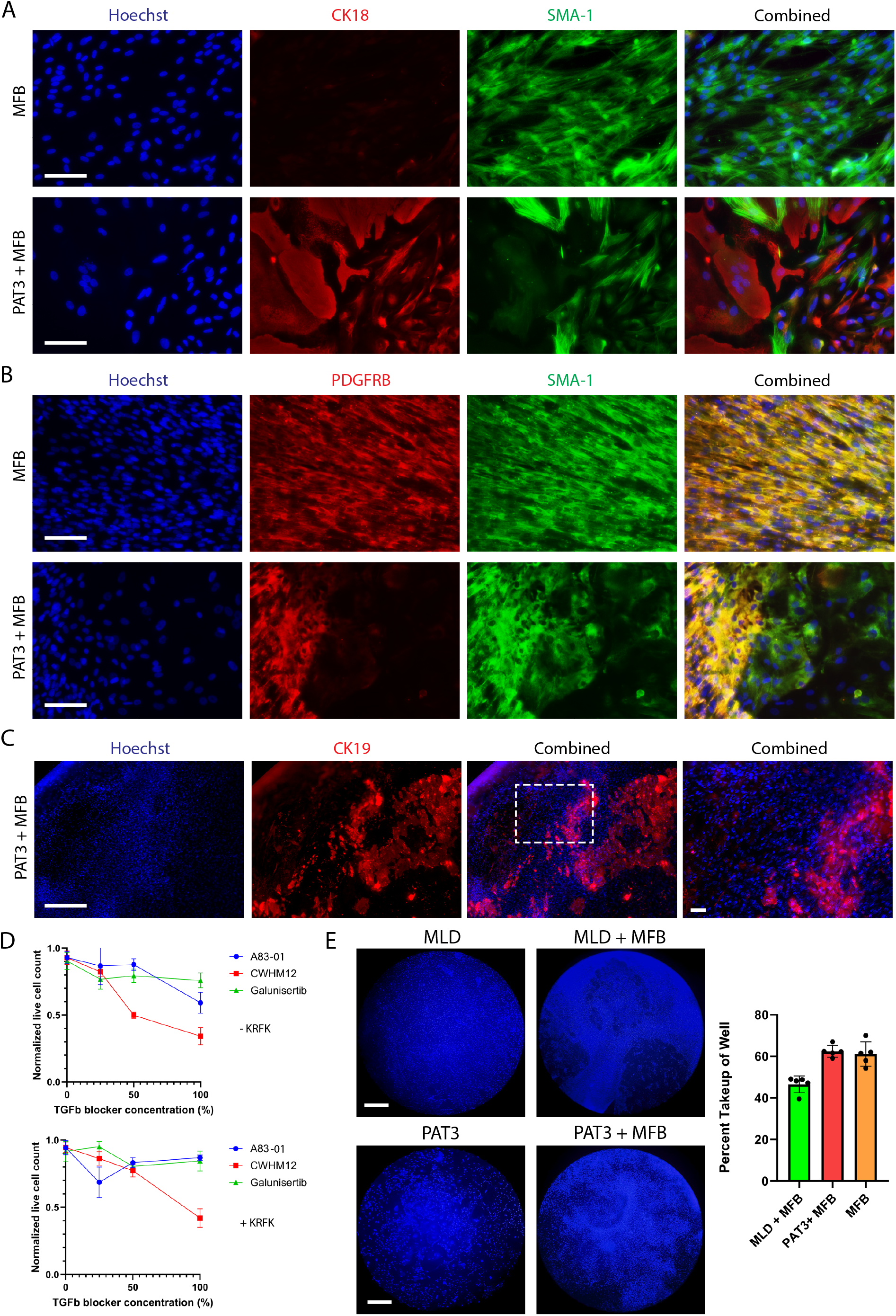
The fibrosis model recapitulates MFB proliferation resulting in formation of monolayers that are predominantly SMA-1+ PDGFR-, and multi-layered spindles that are SMA-1 + PDGFR+. A) MFBs express a-smooth muscle actin (SMA-1) and PAT3s express cytokeratin-18 (CK18). As a control, MFB monoculture is shown in the upper panel. Scale bars, 100 µm. B) Two types of MFB are observed when co-cultured with PAT3. One type only expresses SMA-1 and tends to be monolayered and the other type also expresses platelet derived growth factor receptor beta (PDGFRB) and tends to be part of densely packed spindles. Nuclei not associated with either SMA-1 or PDGFRB belong mostly to the biphenotypic hepatocytes/cholangiocytes of PAT3. A control MFB monoculture is shown in the upper panel. Scale bars,100 µm. C) Low magnification image of a 96 well with a fibrotic scar. MFBs are represented by areas of dense nuclei (Hoechst stain). The organoid cells are labeled with CK19 and partially overlap with the dense MFB areas. Scale bars, 500 µm. D) CWHM12, an antagonist of cell membrane a-integrin mediated TGFP activation, inhibits MFB proliferation with or without the presence of a TGFP inducer, KRFK. Line graphs exclude/include KRFK (top/bottom), and also show other significant TGFP inhibitors A83-01 and Galunisertib (mean ± SD, n=8). E) MFB proliferation is significantly stronger in the presence of PAT3 compared to MLD. Scale bars, 1 mm. The corresponding quantitative percentage of densely populated MFB areas in double co-cultures of organoids is shown. (mean ± SD, n = 5).

**Figure 4:**
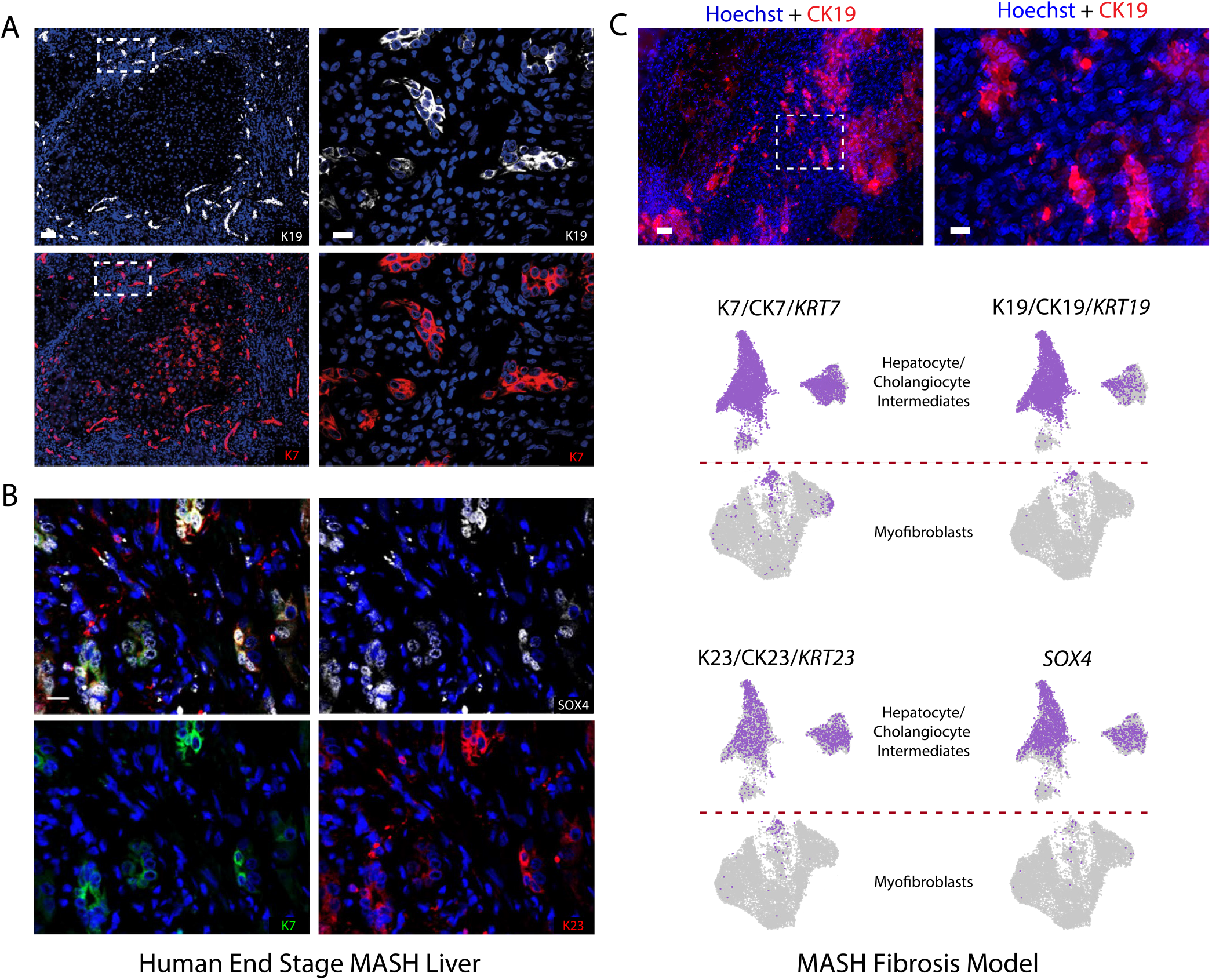
The MASH fibrosis model recapitulates major characteristics of MASH liver fibrotic scars. A, B) Fibrotic areas in end stage MASH patient liver reproduced from IHC staining of end stage MASH liver sections (6). A) Biphenotypic cells associated with high density MFBs labeled with K19/CK19 (upper panel) and K7/CK7 (Lower panel). High density MFBs are represented by high density nuclear areas (blue). B) Biphenotypic cells associated with high density MFBs are labeled with SOX4 (white), K7/CK7 (green) and K23/CK23 (red). High density MFBs are represented by high density nuclear areas (blue). C) Fibrosis model (upper panel). Biphenotypic cells associated with high density MFBs labeled with K19/CK19. The right image highlighted by the white frame shows higher magnification labeling. High density MFBs are represented by high density nuclear areas (blue). Single cell RNAseq shows that biphenotypic cells expressing *KRT19* also express *KRT7, KRT23, SOX4*.

### MASH patient liver-derived organoids enable strong proliferation and scar-like multi-layer assembly of MFBs

During the first ∼5 days after triple co-culture establishment, SMA-1 (*ACTA2*) positive MFBs proliferate strongly and form 3D fibrotic scar-like structures, while organoid-derived cells and MCs do not proliferate. These fibrotic scars exhibit completely novel properties, such as formation of multilayered MFB ‘strings’ surrounding and partially overlaying monolayer clusters of organoid cells strongly reminiscent of fibrotic scars in the human liver (**Figure 3B, C, E**). In contrast, areas of lower MFB density show flattened MFBs (**Figure 3A**). In monoculture, MFBs also exhibit strong proliferation but result in uniformly distributed spindle-like SMA-1+ and PDGFRB+ cells with less scar-like grouping.

Strikingly, only the most condensed areas of multi-layered MFBs contain MFBs that strongly express PDGFRB, a classical marker for activated MFBs, while the less dense areas of MFBs only express SMA-1 (**Figure 3B, C**). The fibrotic scar structures form in ∼5 days and are stable in fibrosis medium (see methods) for at least an additional 10 days. Within this 3D scar-like structure, any organoids cells present are ‘flattened out’ by the MFBs which proliferate on top of the organoids as they are added several days after seeding the organoids.

Interestingly, MFBs can proliferate strongly in the absence of any organoids (**Figures 3A, B upper panels**). This is fully dependent on the presence of the BME (Matrigel) coating on the culture well surface (**Figure S4A**). BME contains approximately 2 ng/ml Transforming Growth Factor beta (TGFβ) and our observation that TGFβ inhibitors significantly reduce MFB proliferation in monocultures (**Figure 3D**) strongly suggests that BME drives MFB replication in monoculture, at least in part through TGFβ.

Healthy organoids significantly inhibit MFB replication and do not allow MFBs to form multi-layered fibrotic scars directly on top of them (**Figure S3**). In contrast, MASH organoids promote MFB replication and allow multi-layered scar tissue formation even if the MFBs are sitting directly on top of the organoids, while the organoids themselves are in contact with the BME coat of the culture well (**Figure 2A, Figure 3B, C, E, Figure S3**). In contrast to MASH patient liver-derived organoid cells, healthy liver organoid cells are not only smaller in size (**Figure 2**) but also fail to immediately stop proliferation in differentiation medium, as they continue to proliferate slowly for a few days. Thus, to achieve a similar organoid cell count after 5 days in culture, healthy liver-derived organoid cells must be added at a 1:2 ratio to MASH liver-derived organoid cells (**Figure S4B**).

### The MASH fibrosis model recapitulates fibrotic scar features of MASH patient liver

The MASH patient liver-derived fibrosis model exhibits structures with high similarity to fibrotic scars of end stage MASH liver (**Figure 4)**. Previously published microscopy images of MASH patient liver slices (6) are strikingly similar to our fibrosis model, especially in the general arrangement of parenchymal cells vs MFBs. In the human MASH liver section, biphenotypic parenchymal cells are labeled via IF microscopy CK7+, CK19+, CK23+ and SOX4+ and in the fibrosis model they are labeled via microscopy CK19+ and via single cell data *KRT7+, KRT19+, KRT23+*, and *SOX4+* (see single cell section below for details) the latter two being classical markers of inflamed MASH hepatocytes (6, 12) (see also **Figure S7**). In both cases, the biphenotypic cells are interspersed within the densely packed fibrotic bands of MFBs (areas of densely packed nuclei) and even form similar small clusters of a few cells each.

### MASH liver-derived organoids promote differentiation of scar associated TREM2+ MPs

Peripheral blood-derived TREM2+ CD68+ activated MPs have emerged as a crucial proinflammatory MP population that strongly expands during human MASH development while Kupffer cell populations diminish (7). These MPs are situated at the boundary of active fibrotic areas and physically interact with MFBs in scar tissue (7). Intriguingly, the same group directly showed that isolated MASH liver-derived TREM2*+* CD68*+* MP supernatant promotes human MFB proliferation at least in part via secreted PDGFB (7). However, unlike in humans, TREM2*+* CD68*+* MPs in mice have recently been implicated in fibrosis resolution and thus assigned an anti-inflammatory function (13).

A striking feature of our fibrosis model is the high physical affinity of peripheral blood MC-derived MPs to MFBs. MPs will preferentially accumulate on MFB-dense areas while avoiding areas with parenchymal organoid cells **(Figure 5A)**. The binding of the MPs to the MFB surface of the scar structure starts around 2-3 days after addition of human peripheral blood-derived MCs and human HSC-derived MFBs onto organoid containing wells (**Figure 1**). Full MC differentiation into MPs is characterized by enlargement, the presence of granules, morphological changes (ranging from flattened, circular cellular bodies to largely branched cellular bodies), and expression of CD68, a classical MP marker, as well as TREM2. Notably, MPs irreversibly bind to MFBs throughout the co-culture’s lifetime, with the longest tested time in this study spanning 15 days of co-culture.

**Figure 5:**
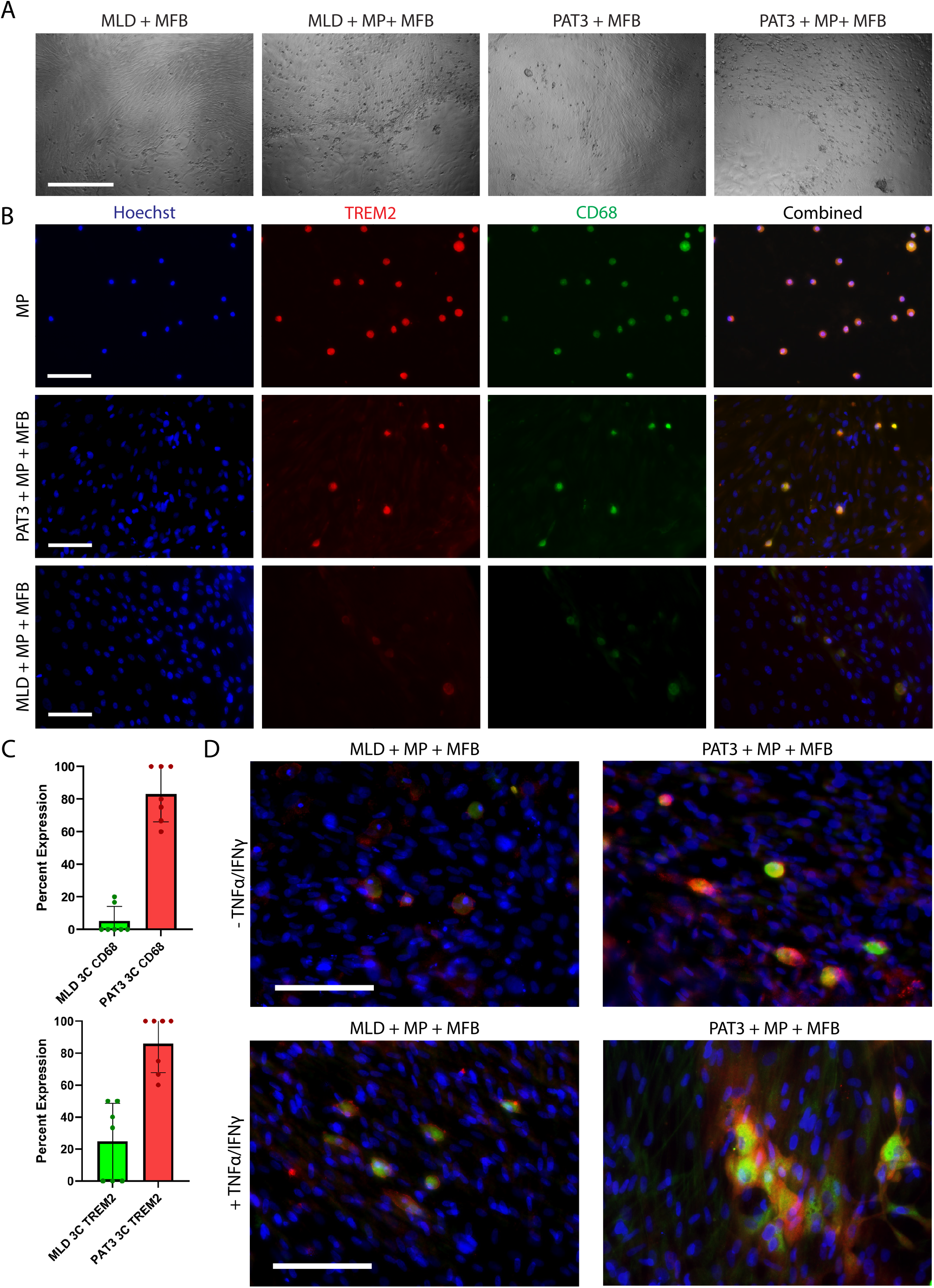
Fully differentiated MPs within the fibrosis model exhibit major characteristics of scar-associated MFBs in the human MASH liver. A) MPs preferentially bind to the surface of MFBs. MFBs form the fibrotic strands in the upper parts of the images and organoid cells form the lower non-fibrotic areas of the images. Scale bar, 500 µm. B) Activated MPs in the context of the fibrosis model are strongly TREM2+ CD68+. Upper panel: MCs in monoculture spontaneously differentiate to TREM2+ CD68+ MPs on BME-coated wells. Middle Panel: MCs bound to MFB similarly differentiate into activated MPs and acquire a TREM2+ CD68+ state in the presence of PAT3. Lower panel: MCs bound to MFB differentiated into activated MPs express TREM2 and CD68 only weakly in the presence of MLD. Scale bar, 100 µm. C) The percentage of TREM2+ CD68+ MPs are substantially increased in PAT3 triple co-cultures than in MLD triple co-cultures (mean± SD, n = 8). Upper panel: CD68. Lower panel: TREM2 D) TNFa and IFNy induce strong branching of pSTING+ CD68+ MPs in the fibrosis model. Left panels: Triple co-cultures containing MLD do feature MPs bound to MFB, but only express low levels of pSTING (red) and CD68 (green) and do not branch after the addition of TNFa and IFNy. Right panels: MPs in triple co-cultures containing PAT3 organoids are pSTING+ CD68+, and flatten and branch extensively in response to TNFa and IFNy. Scale bar, 100 µm.

Both MCs in monoculture and MASH organoid triple co-cultures spontaneously converted to TREM2+ and CD68+ MPs following our culturing procedures (see methods) and remained activated for at least another 10 days. TREM2+ and CD68+ differentiated MCs in monoculture are solely dependent on the presence of BME coating on the culture well surface (**Figure 5B**), similar to monoculture MFB proliferation (**Figure S4A**). Notably, while MC differentiation into MFB bound TREM2+ CD68+ MPs is markedly enhanced in the presence of MASH liver organoids, the presence of healthy liver-derived organoids suppresses MFB bound MPs TREM2 and CD68 expression (**Figure 5B, 5C**). Thus, healthy liver-derived organoids likely produce factors that inhibit the differentiation of MFB-stabilizing TREM2+ CD68+ MPs, similar to their suppressive effects on MFB proliferation (**Figure 3E**). TNFα and IFNγ induced large-scale flattening and branching of MFB-bound pSTING+ CD68+ MPs occurs in the presence of MASH liver organoids, but not healthy liver organoids **(Figure 5D)**. These branched pSTING+ CD68+ MPs are very similar to the proinflammatory fibrotic scar-associated MPs observed previously in human MASH liver (7).

### Single cell sequencing of the MASH organoid fibrosis model reveals senescent biphenotypic cells and activated MFBs

To further assess the relevance of our model for fibrotic scars in MASH patients, we performed single cell sequencing of our 3D fibrosis model with MASH patient liver-derived organoids. As a direct control, we sequenced an identical model grown in parallel with the same MFB and MP batches, but with healthy male donor-derived organoids instead of MASH patient liver-derived organoids. Both models were treated in parallel under identical conditions (see methods). For the last 3 days before single cell harvest, 0.4 mM oleic acid and 0.4 mM palmitic acid were added to the fibrosis medium in an effort to more closely mimic the concentrations of these important free fatty acids present in human MASH liver (14). Both types of triple co-cultures were subjected to single cell preparation using TrypLE, which preserved the parenchymal cells and MFBs, but mostly destroyed MPs due to their higher sensitivity to trypsin digestion compared to the other cell types. Sequencing of both co-cultures was done on the same flow cell, and identical data processing was used for acquiring the RAW data (see methods). Importantly, cell types were then annotated using existing RAW data from the Henderson lab, who performed an extensive single cell analysis of a mixture of 3 MASH and 2 Alcoholic cirrhosis livers at the time of orthotopic liver transplantation as well as a mixture of 5 healthy liver donors as control (7).

**Figure 6A** shows a UMAP containing single cells from both co-cultures. Surprisingly, the only cell population that is highly similar between the two co-cultures is a subtype of MFBs. **Figure 6B** shows annotations of cell types according to the Henderson lab’s single cell data mentioned above. The parenchymal cells in the ‘healthy’ co-culture consist of several distinct types of cholangiocyte or hepatocyte-like cells and one subtype of MFBs. In contrast, the MASH fibrosis model contains only two major subtypes of biphenotypic cholangiocytes/hepatocytes and several subtypes of MFBs. Overall, the fibrosis model aligns closely with the published fibrotic single-cell data, whereas the ‘healthy’ triple co-culture shows substantially weaker alignment with healthy human livers. This likely reflects the fact that organoids-derived from healthy liver tissue predominantly exhibit a cholangiocyte phenotype with limited hepatocyte features, whereas MASH organoids consist of biphenotypic cells that closely resemble those found in human MASH liver. These biphenotypic cells co-express cholangiocyte and hepatocyte markers and likely arise from activated bipotential stem cells, which themselves originate from cholangiocytes in a process called ductular reaction (6).

**Figure 6:**
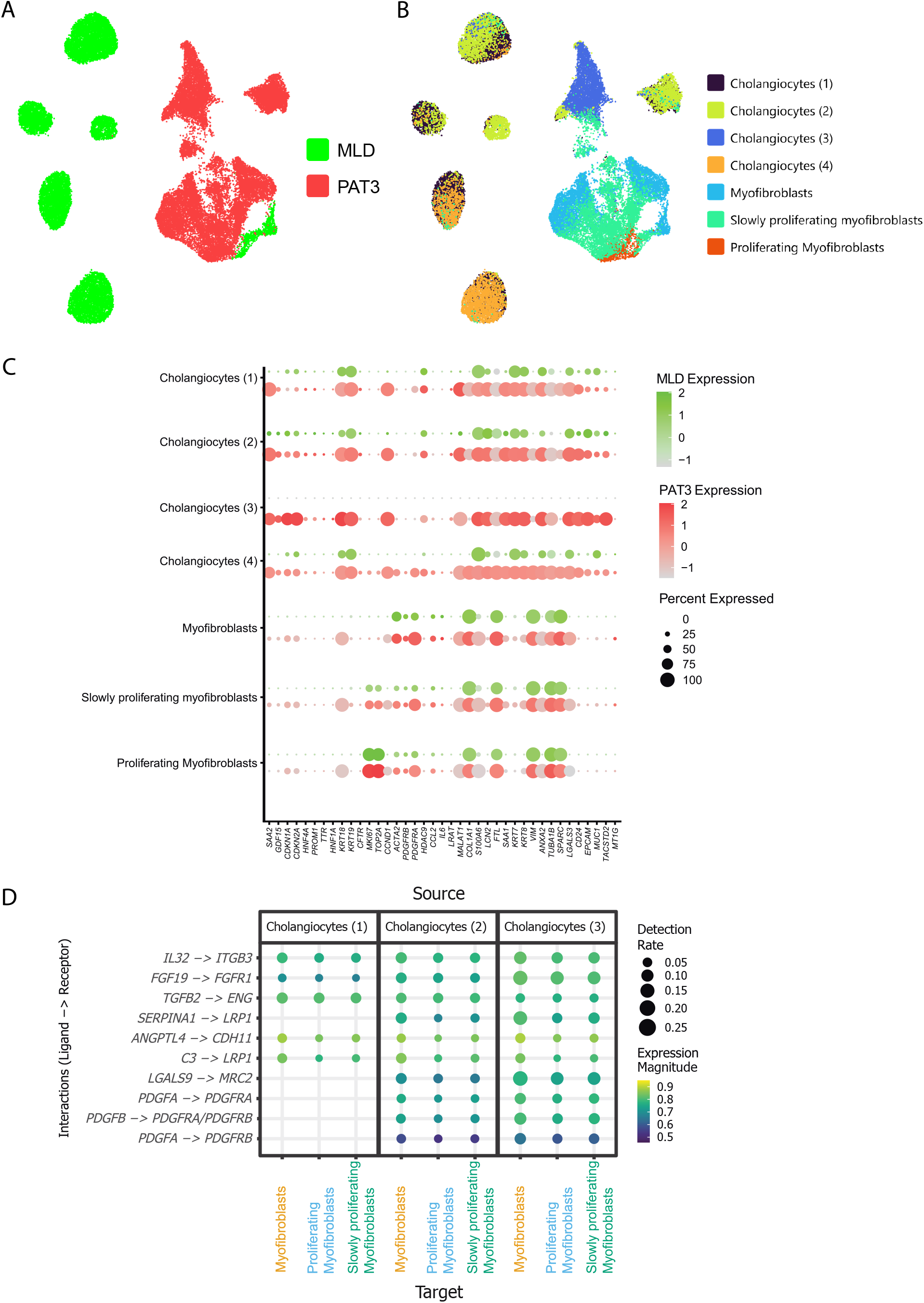
Single Cell analysis of the MASH fibrosis model reveals high similarity with fibrotic scars of human end stage fibrotic liver. A) Combined UMAP of MLD and PAT3 co-cultures showing parenchymal cells and MFBs. Note that there is no meaningful overlap of any MLD and PAT3 populations. MPs are not depicted as the single cell procedure was not compatible with obtaining the majority of intact MPs. B) PAT3s, but not MLDs, strongly stimulate MFB proliferation and activation. Cell types were predicted by unsupervised alignment of the RAW data with published single-cell data from five pooled healthy donor livers and five pooled end stage fibrotic patient livers (10). Both hepatocyte/cholangiocyte and MFB populations are distinct between MLD control co-cultures and PAT3 fibrosis co-cultures. This distinction persists even though the MFBs originate from the same batch and the resulting MLD and PAT3 co-cultures were grown in parallel. C) Gene expression patterns of the predicted cell types of the PAT3 fibrosis model. The dotplot shows expression patterns of both computed and manually curated genes that most strongly discriminate between cell types and are also predicted to be biologically relevant. Note that cholangiocyte type 3, which is the most biphenotypic and the most senescent cell type, only appears in the PAT3 fibrosis model, but not in the MLD control model. Note expression of senescence genes p16 *(CDKN1A)* and p21 *(CDKN2A)* are strongest in cholangicyte subtype 3. The expression pattern of subtype 3 cholangiocytes strongly aligns with that of biphenotypic senescent cells found in human end stage fibrotic MASH liver (10). D) Predicted ligand-receptor interactions between cholangiocyte subtypes and MFB subtypes in the PAT3 fibrosis model. Note that cholangiocyte subtype 3 is the most senescent and most bipotential subtype and importantly, is not present in the MLD control model. The cholangiocyte 3 subtype produces abundant SASP mRNAs in the fibrosis model, such as *IL32, SERPINA1*, and *PDGFNB*. Several of these are predicted to interact with a cognate receptor on the MFBs. The ligand receptor pairs shown here all have been observed and/or predicted in human MASH.

Importantly, in a manner very similar to human biphenotypic cells, the fibrosis model’s cells strongly express many classical senescence markers. These include p16 (*CDKN1A*), p21 (*CDKN2A*), beta galactosidase (*GLB1*), γ*H2AX*, and others (**Figure 6C, Figure S7**). Additionally these cells show strong induction of proinflammatory senescence associated secretory phenotype (SASP) mRNAs, including *SAA1, SAA2, GDF15, CCL2, CCDN1, CXCL2, IL32*, TGFβ (*TGB1* and *TGFB2*) and *PDGFA* (**Figure 6C, Figure S7**), all of which are also well characterized markers of human MASH and increase with severity of the disease (7, 15-17).

Within the senescent biphenotypic cells, two major subpopulations are present, both of which resemble senescent biphenotypic cells in end stage MASH liver. In this cell type, strongly expressed hepatocyte markers include *HNF4A* and *HNF1A* - two key transcription factors that mediate hepatocyte fate - as well as *TTR* and *SERPINA1*, which encode major secreted hepatocyte proteins and glutamine synthetase (*GLUL*). We see all these genes expressed (**Figure 6B, C, Figure S7**). Biphenotypic cells from the human MASH liver exhibit strongly reduced expression of albumin *(ALB), CYP3A4*, and other classical hepatocyte mRNAs, a pattern that we similarly observed in the biphenotypic cells of our fibrosis model. Our healthy co-culture system lacks true hepatocytes and instead predominantly contains the three cholangiocyte subtypes predicted by the Henderson lab’s single-cell study and therefore cannot accurately model hepatocytes in healthy human liver.

Remarkably, there are ∼10 fold more MFBs in the MASH co-culture than in the healthy co-culture, (**Figure 6A and B**) broadly consistent with our functional data that show a less pronounced but still statistically significant difference (**Figure 3E**). This suggests highly significant inhibition of MFB replication specifically in the presence of healthy organoids **(Figure 3)**. However, the true abundance of each cell type may be partially obscured by their differential sensitivity to the single-cell dissociation procedure. Indeed, tightly packed MFBs may be more resistant to trypsin digestion than MFB monolayers. Our single-cell analysis reveals that patient liver-derived organoids are markedly more profibrotic than those-derived from healthy liver and strongly support both MFB proliferation and overexpression of profibrotic genes *PDGFRA*, SMA-1 *(ACTA2), COL1A2, COL3A1*, and *DCN* (**Figure 6C, Figure S7**). Furthermore, MFBs consist of non-replicating, slow, and high replicating types. MFBs also express TGFβ1 (*TGFB1*), which is likely part of an autocrine loop to support MFB growth (**Figure S7)**. A subset also expresses PDGFRB, which is consistent with our earlier data (**Figures 2, 3, and 7**). Interestingly, MASH organoid biphenotypic cells, but not healthy organoid cells, produce *TGFB2*, which also is known to induce MFB replication (**Figure S7**). This cell type specific difference between *TGFB1* and *TGFB2* expression is not found in mice, supporting the utility of using a human 3D system.

**Figure 7:**
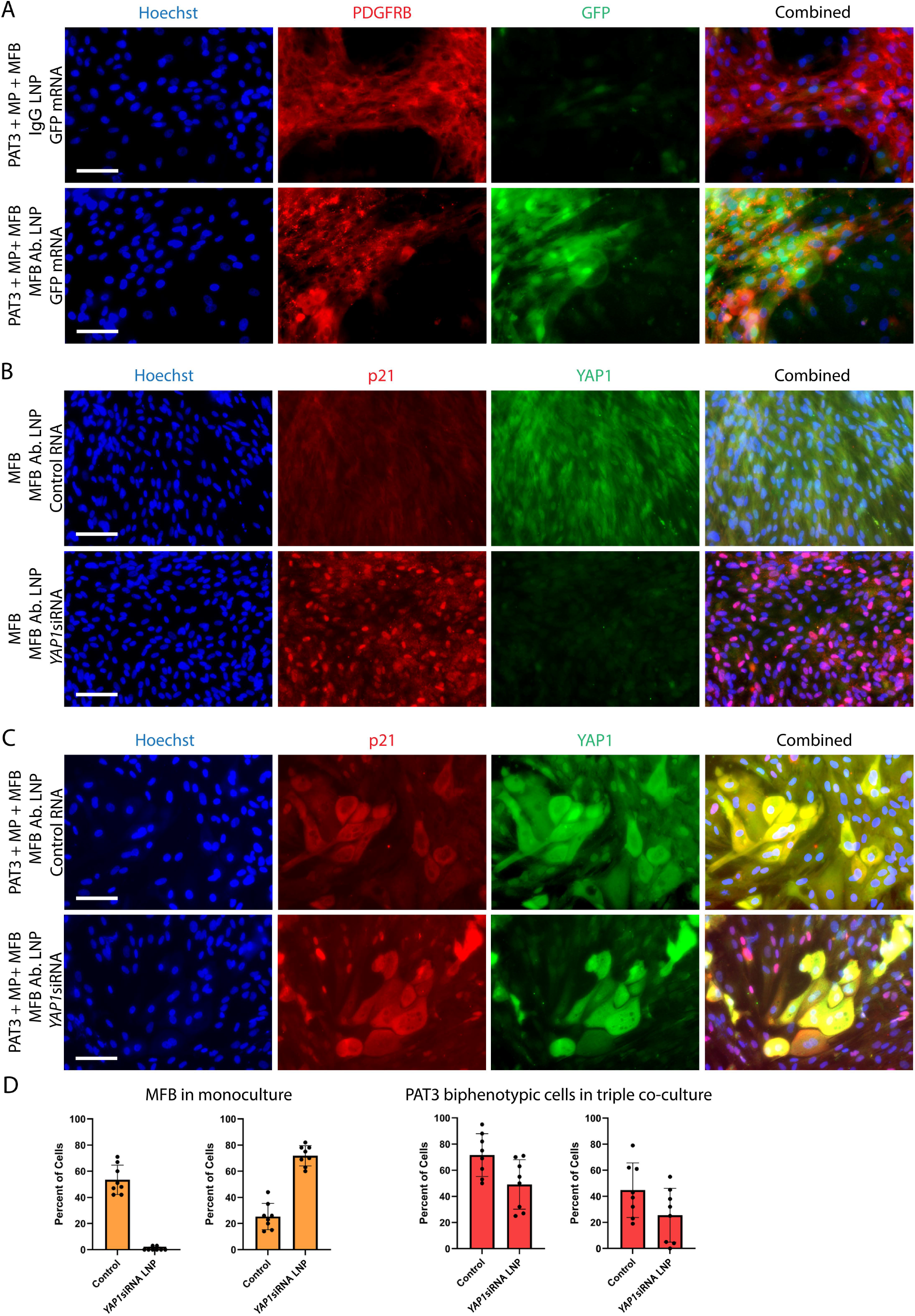
LNPs specific for MFBs that are formulated with YAP1siRNA knockdown *YAP1* only in the MFBs, but not the biphenotypic hepatocytes/cholangiocytes of the fibrosis model. A) Highly efficient MFB specific LNP-mediated targeting in the fibrosis model. Upper panel: IgG control LNP formulated with GFP mRNA in the context of lower density MFBs allows relatively weak, yet specific targeting. Lower panel: An LNP conjugated with an anti-MFB specific antibody allows for highly efficient uptake and expression of GFP mRNA in MFBs, but not PAT3 biphenotypic organoid cells. Scale bar, 100 µm. B) While IgG control LNP shows no change in p21 and YAP1 expression, the YAP1siRNA LNP not only causes almost complete knockdown of nuclear YAP1 in MFBs, but also results in nuclear overexpression of the senescence marker p21. Scale bar, 100 µm. C) YAP1siRNA LNP addition to fibrosis triple co-cultures induces MFB specific gene editing. No change in either YAP1 or p21 is seen in biphenotypic cells, while *YAP1* knockdown and p21 overexpression are seen in MFB. Note that PAT3 biphenotypic cells express large amounts of YAP1 and p21 mostly in the cytoplasm under all conditions. No change in either YAP1 or p21 is seen in hepatocytes. Scale bar, 100 µm. D) Quantitative analysis of nuclear YAP1 and nuclear p21 in MFBs from B), and from biphenotypic cells from PAT3 triple co-cultures in C). Left graphs: Nuclear YAP1 protein expression. Right graphs: Nuclear p21 protein expression (mean ± SD, n = 8).

**Figure 6D** shows prominent ligand receptor pairs. Ligands are expressed on the cell surface and/ or secreted as part of the SASP by the senescent biphenotypic cells and receptors are located on the cell membrane of the MFBs in the MASH patient model, but importantly, not in the corresponding healthy model. We identified 11 ligand receptor pairs. The best previously documented of these are the *PDGFA/B-PDGFRA/B* pairs (7) with less evidence for *TGFB2-ENG, FGF19-FGFR1, SERPINA1-LRP1, IL32-ITGB*, and *LGALS9-MRC2*. Overall, this represents strong supporting evidence that the SASP of the senescent MASH organoids is responsible for the strong MFB proliferation on top of the organoid cells, a location that does not allow direct physical contact with the latent TGFβ complex rich BME coating of the well. **Figure S7** shows UMAPs of the fibrosis model that show cell type specific mRNAs, including those involved in senescence, SASP, proliferation, and extracellular matrix production. Taken together, our signal cell data confirm the close resemblance of our human fibrosis model to the MASH liver fibrotic scar.

### LNP-mediated gene silencing of *YAP1* induces senescence specifically in MFBs

To assess the compatibility of our 3D MASH fibrosis model with gene silencing techniques, we chose to test several types of LNPs that were designed to specifically target MFBs within our co-cultures. Inhibiting the proliferation of MFBs is an important therapeutic goal of treating MASH (18), but any such method must also avoid targeting other cell types to prevent unwanted, off-target side-effects. Two important cell types that have historically been difficult to avoid targeting are regenerative hepatocytes and bipotential stem cells, both of which are present within the regenerative nodules of human cirrhotic MASH liver and surrounded by fibrotic scar tissue (6). First, we tested an LNP with the 4A3-sc8 ionizable lipid (Echelon Biosciences) and formulated with GFP mRNA. This ligand has previously been shown to facilitate highly efficient LNP delivery of nucleic acids into dividing cells (11). This 4A3 LNP shows considerable specificity for a subset of MFBs, but no other cell types in the fibrosis model although it does not allow very high expression of GFP mRNA. (**Figure 7A upper panel**). We then tested the same 4A3 LNP conjugated with an MFB specific antibody showing significantly improved efficiency of GFP mRNA expression with very high specificity for MFBs (**Figure 6A, lower panel)**.

An isotype control LNP showed only low levels of GFP mRNA expression similar to the 4A3 LNP (not shown). 4A3LNP conjugated to a MFB specific antibody resulted in strongly improved MFB specific GFP expression **(Figure 7A)**.

To induce senescence in proliferating MFBs, an important therapeutic goal in MASH (18), we chose to silence *YAP1* expression in MFBs. Indeed, previously published data have shown that liver MFB proliferation is at least partially dependent on nuclear YAP1 expression (18). To achieve *YAP1* silencing, we formulated MFB specific antibody conjugated LNPs with anti-*YAP1* siRNA. We demonstrated reduction in nuclear YAP1 mRNA in MFB monocultures by > 95% (**Figure 7B, D**). In triple co-culture organoids treated with MFB specific antibody conjugated LNPs formulated with anti-*YAP1* siRNA, a similarly strong *YAP1* silencing was seen specifically in MFBs, but not in hepatocyte/cholangiocytes (**Figure 7C, D**), indicating high cell type specificity. *YAP1* silencing also induced strong nuclear induction of the senescence marker p21 in the MFBs, but not in the hepatocyte/cholangiocyte organoid cells, as shown by double IF microscopy of YAP1 and p21 (**Figure 7B, C, D**).

## Discussion

For this work we selected one ‘average’ male end stage MASH organoid line (PAT3) and one ‘average’ healthy male living donor line (MLD) from our previously published 6 end stage MASH patient liver-derived organoid lines and 3 healthy donor liver-derived organoid lines (3). The criteria for selection were average growth speed during expansion and average senescence marker over expression after full differentiation. We are therefore confident that our present data reasonably reflects ‘average’ human end stage MASH.

We find that combining MASH patient liver-derived organoids with primary human HSCs and human peripheral blood MCs causes rapid spontaneous assembly of 3D MASH fibrotic scar-like structures driven by strong MFB proliferation. Single cell sequencing shows that these triple co-cultures contain strongly senescent biphenotypic hepatocyte/cholangiocyte intermediates that over express secreted factors capable of stimulating MFB proliferation. Additionally, MCs differentiate into scar-associated TREM2+ MPs that are likely proinflammatory as previously described for human MASH liver (7). We also established LNP-mediated knockdown of nuclear YAP1 specifically in MFBs, causing them to become senescent and non-proliferative while not affecting other cell types.

Our findings suggest that the epigenetic memory of the human MASH liver persists in the corresponding organoids within the context of our fibrosis model. Indeed, the organoids are only 4-6 passages removed from the actual MASH liver tissue and unlike iPSC-derived hepatic organoids are not subjected to any genetic reprogramming. Therefore, the MASH liver-derived fibrotic system can reflect pathological pathways that have accumulated in MASH patient liver over the entire time of its development, which spans 10-20+ years.

The prominence of senescent hepatocyte/cholangiocyte biphenotypic cells in our fibrosis model replicates a profound feature in human MASH liver, which has been poorly replicated in mice and in iPSC hepatic organoid models. The biphenotypic cells simultaneously overexpress the classical senescence markers p16 (*CDKN1A*) and p21 (*CDKN2A*) (6, 7), and exhibit a senescence associated secretory phenotype (SASP), which includes the cytokines *TGFB1* (log2>2), *GDF15* (log2 > 2.5), *IL32* (Log2>5), *CCL2* (log2>2), the growth factor PDGFA (log2>2), and serum amyloids *SAA1* (log2>5) and *SAA2* (log2>5), all of which are profoundly proinflammatory and attract immune cells such as MCs and MPs. All these SASP components have been previously described to increase with the severity of MASH liver fibrosis (18-26).

Single cell sequencing further revealed several new ligand receptor pairs between senescent biphenotypic cells and MFB receptors that could explain the strong MFB replication. Among those are *IL32-ITGB3, FGF19-FGFR1, SERPINA1-LRP1*, while *PDGFA/B-PDGFRA/B* was expected based on several previous publications (7, 27). Importantly, blood concentrations of all the above ligands have been reported to positively correlate with MASH severity (28-31). All these ligands are senescent biphenotypic cell-secreted proinflammatory proteins that are part of a SASP, similar to the SASP found in human MASH liver (7).

We confirmed an important previously functionally described human MASH pathway, namely the differentiation of proinflammatory TREM2+ CD68*+* scar-associated MPs from peripheral MCs (7). It is worth noting that mouse models of TREM2*+* MPs do not reflect the proinflammatory, human TREM2+ MPs, but instead exhibit anti-inflammatory TREM2+ MPs associated with both lipid-laden hepatocytes and fibrotic scars (11). We show that the TREM2+ CD68+ MPs bind very strongly to the MFBs. Since our single cell analysis shows that the MFBs in the MASH 3D model, but not the healthy control model express large amounts of collagens (*COL1A1, COL1A2, COL3A1* and others; log2 fold change for all > 5), it is conceivable that MFB-secreted extracellular matrix coating on the cell surface directs MPs toward the TREM2+ fate, similar to the BME-coated wells with MC monocultures containing collagen. It will be interesting to investigate the exact mechanism(s) of *TREM2* inhibition in the healthy control model for therapeutic purposes.

Our use of LNPs has several advantages over other types of genetic manipulation of 3D organoid systems, such as speed, ease of application, and the ability to target non-dividing cells. Indeed, we show that MFB specific antibody conjugated LNPs formulated with anti-*YAP1* siRNA strongly knock down YAP1 protein and induce p21 mediated senescence in MFBs while not affecting other cell types.

MFB senescence is an important therapeutic goal because it is an important first step towards fibrosis resolution. Further, senescent cells become highly sensitive to senolytics. We are currently exploring the possibility to induce senolytic cell death in senescent MFBs by simultaneously inducing senescence and also knocking down genes that prevent cell death specifically in senescent cells. Killing senescent MFBs with a one-two punch might be a promising strategy to alleviate advanced MASH. Importantly, such senolytic killing should be safe because of our use of cell type specific knockdown which could prevent unwanted side effects.

In a broader context, our model is compatible with high throughput screening techniques, as our data were all collected in 96 well plates, and the time from frozen cell stock to fully differentiated model is only 2-3 weeks, shorter than other classical MASH models. This opens novel avenues to screen for improved anti-fibrotic therapeutics, both small molecule and LNP-mediated.

Our MASH fibrosis model has some limitations. Its parenchymal cells are mostly senescent biphenotypic cells, rather than more ‘pure’ hepatocytes, and thus the system seems limited to observe interactions within a fibrotic scar itself rather than within areas of human MASH liver that lack MFBs. Most prominently, our system lacks endothelial cells and associated blood circulation. However, our model could be further augmented by adding liver sinusoidal endothelial cells, which have important functions in the MASH fibrotic scar and also present a major barrier to the entry of LNPs and peripheral MCs *in vivo*. In addition, it might be possible to transplant the fibrosis model into immune compatible mice to study its integration into the host’s blood circulation.

## STAR Methods

| REAGENT or RESOURCE | SOURCE | IDENTIFIER |
| --- | --- | --- |
| **Antibodies** |  |  |
| yap1 | proteintech | Cat# 66900-1-Ig; RRID:AB_2882229 |
| CD68 | Thermo Fisher (invitrogen) | Cat# 14-0688-82; RRID:AB_11151139 |
| α-sma1 (ACTA2) | R&D Systems | Cat# MAB1420; RRID:AB_262054 |
| Cytokeratin 18 (CK18) | proteintech | Cat# 10830-1-AP; RRID:AB_2133164 |
| Cytokeratin 19 (CK19) | proteintech | Cat# 10712-1-AP; RRID:AB_2133325 |
| PDGFRβ | biotechne (R&D Systems) | Cat# AF385; RRID:AB_355339 |
| TREM2 | ThermoFisher | Cat# PA518763; RRID:AB_10984816 |
| phospho-STING (Ser366) (E9A9K) | Cell Signaling Technology | Cat# 50907; RRID:AB_2827656 |
| CDNK1A/p21 | ABclonal | Cat# A19094; RRID:AB_2862586 |
| Anti-Rabbit IgG (secondary, 488) | Cell Signaling Technology | Cat# 4412; RRID:AB_1904025 |
| Anti-Mouse IgG (secondary, 488) | Thermo Fisher (invitrogen) | Cat# A28175; RRID:AB_2536161 |
| Anti-Mouse IgG (secondary, 546) | Thermo Fisher (invitrogen) | Cat# A11003; RRID:AB_2534071 |
| Anti-Goat IgG (secondary, 568) | Thermo Fisher (invitrogen) | Cat# A11057; RRID:AB_2534104 |
| Cy3-Conjugated Anti-Rabbit (secondary, 550-570) | proteintech | Cat# SA00009-2; RRID:AB_2890957 |
| **Fluorescent Dyes** |  |  |
| Hoechst 33258 Solution | Sigma | Cat# 94403-1ML; RRID:AB_2651133 |
| HCS LipidTOX (Red Neutral Lipid Stain) | invitrogen | Cat# H34476 |
| **Chemicals, Peptides, and Recombinant Proteins** |  |  |
| IFN-γ | EMD Millipore Corporation | Cat# GF305 |
| BMP-7 | Peptotech | Cat# 120-03P-50UG |
| TNF-α | Millipore Sigma | Cat# SRP3177-50UG |
| Recombinant Human HGF | Peptotech | Cat# 100-39-50UG |
| Advanced DMEM/F12 | Thermo Fisher | Cat# 12634010 |
| HEPES (1M) | Thermo Fisher | Cat# 15630-080 |
| GlutaMAX | Thermo Fisher | Cat# 35050-061 |
| Penicillin-Streptomycin | Thermo Fisher | Cat# 15140-122 |
| R-spondin 1 (RSPO1) | Made as in (5) | N/A |
| B27 supplement (w/ Vit A) | GIBCO | Cat# A3582801 |
| B27 supplement (w/o Vit A) | GIBCO | Cat# 12587010 |
| A83-01 | Tocris | Cat# 909910-43-6 |
| N-Acetylcysteine | Sigma | Cat# A9165-25G |
| N2 Supplement | Thermo Fisher | Cat# 17502048 |
| Nicotinamide | Sigma | Cat# N0636 |
| Recombinant Human EGF | Peptotech | Cat# AF-100-15 |
| Gastrin | Sigma-Aldrich | Cat# G9145 |
| DAPT | Sigma-Aldrich | Cat# D5942 |
| Dexamethasone | Sigma-Aldrich | Cat# D4902 |
| Forskolin | Tocris Bioscience | Cat# 2939 |
| ROCK inhibitor (Y-27632) | Abmole | Cat# Y-27632 |
| FGF10 | Peptotech | Cat# 100-26-100UG |
| FGF19 | Peptotech | Cat# 100-32-50UG |
| Noggin | Peptotech | Cat# 120-10C-5UG |
| TrypLE Express | GIBCO | Cat# 12605010 |
| Galunisertib | MedChem Express | Cat# HY-13226 |
| CWHM-12 | MedChem Express | Cat# HY-18644 |
| KRFLK (TFA) | MedChem Express | Cat# HY-P3970A |
| XMU MP-1 | Bio-Techne | Cat# 10-1-6482 |
| BSA, FRACTION. V, Fatty Acid, Nuc, Prot-free | Millipore Sigma | Cat# 126609-5GM |
| RPMI 1640 (1x) | Corning | Cat# 10-040-CM |
| Triton X-100 | Roche Diagnostics | Cat# 10-789-704-001 |
| Cultrex RGF Basement Membrane Extract (BME), Type 2 | Bio-Techne | Cat# 3533-005-02 |
| Human Serum Type AB (MALE) | Millipore Sigma | Cat# H4522-100ML |
| 1% FBS | Life Technologies | Cat# 26010066 |
| Earle's Balanced Salt Solution (EBSS) (Ca<sup>2+</sup>/Mg<sup>2+</sup>) | Thermo Fisher Scientific HyClone | Cat# 12349752 |
| Collagenase D | Roche | Cat# 1108866001 |
| DNaseI | Sigma-Aldrich | Cat# DN25 |
| Wnt3a-Conditioned Medium | Made as in (5) | N/A |
| Bovine Serum Albumin (BSA) | Sigma-Aldrich | Cat# A8806-1G |
| PBS-1x | Life Technologies | Cat# 14190-094 |
| DMSO | Sigma-Aldrich | Cat# D8418 |
| **Oligonucleotides** |  |  |
| GFPmRNA | Weissman lab, University of Pennsylvania |  |
| ON-TARGETplus Human YAP1 (10413) siRNA | Dharmacon | Cat# J-012200-08-0010 |
| ON-TARGETplus Non-targeting Pool | Dharmacon | Cat# D-001810-10-05 |
| **Deposited data** |  |  |
| scRNA-seq data | This study | N/A |

| Cell Sources |  |  |
| --- | --- | --- |
| Patient-derived liver organoid lines | This study | N/A |
| Hepatic Stellate Cells (HSCs) | Sciencell | Cat# 5300 |
| Human Peripheral Blood Monocytes | Perelman Human Immunology Core | N/A |
| Software and algorithms |  |  |
| Loupe Browser | 10x Genomics | https://www.10xgenomics.com/products/loupe-browser/downloads/eula |
| Microsoft Office Suite | Microscotf |  |
| Adobe Creative Cloud | Adobe | https://www.adobe.com/gr_en/creativecloud.html |
| GraphPad Prism (version 9.4.2) | GraphPad | https://www.graphpad.com/ |
| BioRender | BioRender | https://www.biorender.com/ |
| NIS Elements |  |  |
| Other |  |  |
| Nikon Eclipse TE2000-U Inverted Fluorescence Microscope | Nikon |  |

### Human tissues for organoid cultures

All experiments using human tissues were approved by the University of Pennsylvania’s Institutional Review Board in accordance with all relevant ethical guidelines (Penn IRB protocol #828085). Details of all biopsy related procedures were already published in (3). Briefly, wedge biopsies from male donors and male MASH explanted livers were obtained from the Penn Transplant Institute at the University of Pennsylvania. Wedge samples for both types were taken from the peripheral liver edge to ensure a comparable location. In the case of patients withdrawing consent, all tissues and-derived materials from said patient will be discarded.

### Organoid derivation and culture

Isolation both healthy (MLD, male living donor) and diseased liver samples (PAT3, called NASH2 patient in the original publication) we described previously in detail (3). Briefly, Wedge biopsies of liver samples were stored at 4°C in organoid Basal medium (Advanced DMEM/F12, 1x Pen/Strep, 1x GlutaMAX, 1x Hepes) for a maximum of 20 hours. Tissues were then minced with a sterile razor blade and washed with organoid wash medium (DMEM high glucose, 1x GlutaMAX, 1x pyruvate, 1x pen/strep and 1% FBS). Minced samples were placed in digestion solution (EBSS, Collagenase D 2.5 mg/mL, DNaseI 0.1 mg/mL) for 15-30 minutes for healthy donor tissue or 60-90 minutes for diseased patient tissue. MASH liver samples were passed through a 70 µm cell strainer (Thomas Scientific, #410-0002-OEM) and healthy liver samples were unfiltered. Samples were collected through centrifugation at 1400 rpm for 3 minutes and embedded and resuspended in Cultrex Basement Membrane Extract (BME) type 2 PathClear (Milipore-SIGMA, #3532-001-02). Cells were then plated in 50-60 μl-volume of BME onto 24 well suspension plates (Greiner Bio, 662102) and allowed to solidify through 10-15 minutes of incubation at 37°C with 5% CO2. Finally, tissue droplets were overlaid with isolation medium (Basal media, 30% Wnt3a conditioned medium, 25 ng/mL Noggin, 10 µM ROCK inhibitor, 1x B-27 supplement without vitamin A, 1 mM NAC, 10 mM Nicotinamide, 10% R-spondin conditioned medium, 1x N2 supplement, 100 ng/mL FGF-10, 25 ng/mL HGF, 25 ng/mL human EGF, 10 nM leu15 Gastrin, 10 µM Forskolin, and 5 µM A83-01).

The passaging procedure of healthy and diseased liver samples/cells is identical. Established 3D cultures were mechanically sheared in-well by pipetting up and down approximately 15 times with a 200 µl pipette tip (Corning, 4711) mounted on a 1 ml pipette tip (Corning, 9032), yielding a mix of single cells and clumps of approximately 10-50 cells. Cells were suspended into approximately 3 ml of 4°C Basal medium per 50 µL of matrix and left on ice for 5-10 minutes to completely dissolve the BME. After centrifugation at 1400 rpm for 3 minutes and removal of supernatant, the cells were suspended and embedded in 50-60 µL volume droplets containing approximately 2000 cells per droplet, plated on a 24 well plate, and allowed to incubate at 37°C for 10-15 minutes. Once the droplet solidified, the well was overlaid with pre-warmed expansion medium (Basal media, 1x B27 supplement without vitamin A, 1 mM NAC, 10 mM Nicotinamide, 10% R-spondin conditioned medium, 1x N2 supplement, 100 ng/mL FGF-10, 25 ng/mL HGF, 25 ng/mL human EGF, 10 nM leu15 Gastrin, 10 µM Forskolin, 5 µM A83-01).

For optimal expansion of hepatocyte organoids, cell counts per matrix droplets should stay between 200,000 and 1.5 million cells. The approximate splitting ratio and timeline for organoids was 1:4 every 4-8 days, depending on the organoid line. When starting cells from frozen vials, an additional 10 µM ROCK inhibitor Y-27632 added to the expansion medium is necessary for cell survival up to 3 days in culture. Routine splitting does not require ROCK inhibitor Y-27632 as the larger cell chunks increase cell survivability. Assessment of cell viability of these tissues was performed in a previously published experiment (3).

### Primary MFB Culture

Primary human HSCs from healthy donor livers were obtained from Sciencell (cat.# 5300) at passage 1. To simultaneously proliferate and differentiate HSCs into MFBs, a 24 well plate was incubated for at least 2 hours up to overnight with a coating media (Basal media, 1% BME), creating a layer of BME on the bottom of the well. To determine the optimal simultaneous growth and differentiation medium for HSCs, several candidate formulations of media pairings were tested on early passage HSCs. The final fibrosis media yielded the best combined results, consisting of a 1:1 mixture of organoid differentiation media (see below) and RPMI, supplemented with 0.5x Pen/Strep, 0.5x Hepes, 0.5x GlutaMAX, and 1% Human Serum. HSCs were then added directly to the coated wells with this fibrosis media. HSCs overlaid with this media fully differentiated into MFBs over 5-7 days and continued to grow indefinitely.

Due to the cell density and muscle-like properties of the MFBs after proliferation, an additional treatment of 1x PBS -/- alongside TrypLE dissociation was required before routine passaging. First, media in MFB wells of a 24 well plate were replaced with 1x PBS -/- and allowed to incubate for approximately 15 minutes at 37°C to begin detaching the cell sheet from the plate. Afterwards, the 1x PBS -/- was removed and the cells were treated with TrypLE for 4-6 minutes at room temperature and resuspended, resulting in a near-single-cell solution (clumps of approximately 5-10 cells). The cells were then added to Basal media and centrifuged at 1400 rpm for 3 minutes. The supernatant was then removed, cells suspended in fibrosis media and added directly to a BME-coated 24 well plate. For optimal outgrowth of MFBs, a coated well should contain between 50, 000 and 3 million cells, and the approximate splitting ratio and timeline for these organoids is approximately 1:24 every week. ROCK apoptosis inhibitors are not necessary when starting HSCs from frozen vials.

### Human peripheral blood MC harvest

Isolation of primary peripheral blood MCs was done by the Perelman Human Immunology Core, where they performed negative selection on healthy donor blood using the RosettaSep Kit from STEMCELL Technologies. The donor origin of MCs was kept consistent across an experiment. MCs were added to Basal media and spun down at 1400 rpm for 3 minutes. Afterwards, the supernatant was removed, and the cells were resuspended in freezing media (see below) and frozen down at -80°C. MCs did not significantly proliferate within the scope of this experiment and were only used from frozen vials to add onto all-cultures. MCs were cultured in mono, double, or triple co-cultures in the presence of fibrosis media on 1% BME coated wells.

### Freezing and thawing cells

The procedure for freezing down organoid cells and MFBs was identical. Hepatic organoid cells or MFBs were processed by up and down pipetting to produce single cells or small cell chunks. Cells were centrifuged at 1600 rpm for 3 minutes, supernatant was removed, and freezing media (Fetal Calf Serum, 40% DMEM, 10% DMSO) were added to the pellet, resuspended and frozen down at -80°C.

After thawing any cell type, cells in freezing mix were diluted at least 10 times with cold basal media and centrifuged at 1400 rpm for 3 minutes. The supernatant was removed before proceeding with any plating procedures.

### Triple co-culture for fibrosis modeling

A timeline visualizing the establishment of all monocultures and triple co-cultures for fibrosis modeling is provided in **Figure 1**.

Monoculture plating procedures of healthy and diseased liver organoid cells were identical. Monocultures were established on 96 well plates (Greiner Bio, 655090) that had been incubated at 37°C with coating media for a minimum of 1 hour and up to overnight in each desired well. Fully grown organoids in basement membrane extract (BME) droplets on expansion 24 well plates were mechanically sheared in-well by pipetting up and down 5-7 times with a 200 µl pipette tip mounted on a 1 ml pipette tip, yielding cell clumps of 50-200 cells. The cell suspension was removed from the well and placed into approximately 3 ml of 4°C basal medium per 50 µL of BME and left on ice for 5-10 minutes to dissolve the BME. After centrifugation at 1400 rpm for 3 minutes and removal of supernatant, the cells were suspended in organoid differentiation media (Basal media, 1x B-27 supplement with vitamin A, 1x N2 supplement, 1mM N-acetylcysteine, 100 ng/mL FGF-19, 50 ng/mL human EGF, 25 ng/mL HGF, 10nM leu15 gastrin, 25 ng/mL BMP-7, 10 µM DAPT, 3 µM dexamethasone, and 500 nM A83-01) and plated onto a BME pre-coated 96 well plate.

The optimal starting ratio between healthy and diseased hepatic organoids for equivalent monoculture development/differentiation is 1:2. This was because healthy donor cells would continue to grow for a few days after being overlaid with organoid differentiation media while diseased patient cells would immediately stop proliferation. Following this ratio, 1 fully grown well of healthy organoids or 2 fully grown wells of diseased organoids should be separately split into 24 wells of a coated 96 well plate. Over a 10-12 day differentiation period, wells containing healthy and diseased organoids reached equal densities and were fully differentiated.

Monocultures of MFBs or MPs were started from frozen vials as MPs do not have a proliferation period before differentiation and MFBs proliferate excessively if taken from an actively expanding well. In both cases, cells were taken from frozen vials and added to basal media, centrifuged at 1200 rpm for 3 minutes, the supernatant removed, suspended in fibrosis medium, and plated onto a pre-coated 96 well plate. The approximate starting cell density per well for MFBs and MCs for monoculture establishment is ∼15,000 and ∼30,000 respectively. MFBs continued to proliferate and further differentiate while MCs differentiated into MPs over a 5-7 day period. Neither MFBs nor MPs required addition of any factors to differentiate during this period; the 1% BME coating contains various factors like TGFβ that allow for full differentiation of MFBs and MCs into *SMA-1*+ MFBs and *TREM2*+ MPs, respectively.

For double and triple co-cultures, organoid cells were first established as mostly 2D monolayers by mechanical disruption of 24 well BME drops and seeding on BME coated 96 well plates in differentiation medium. After 5-7 days of differentiation MFBs and/or MPs were added according to the above monoculture procedures. Briefly, MFBs and/or MPs from frozen vials were resuspended in fibrosis media and overlaid onto existing 2D organoid cell monocultures and allowed to differentiate over the remaining culture time. Fibrosis media allowed for continued culture of differentiated organoid cells while maintaining optimal conditions for differentiation of MCs and proliferation of MFBs. Triple co-cultures fully differentiated 5-7 days after the addition of MCs and MFBs. Double co-cultures excluding organoid cells but incorporating MCs and MFBs were treated the same way and in parallel on the same plate together with triple-cultures.

Lipid nanoparticles were added at 5 µg/ml final concentration directly to the culture medium 2-3 days before the end of the experiment. Small molecule inhibitors were applied to all applicable cultures 2-3 days before the end of the experiment. At the end of any experiment, cells were fixed with 10% formalin for 10 minutes at room temperature, followed by 0.05% Triton X permeabilization for 5 minutes at room temperature, ending with albumin blocking with 0.1% BSA in PBS -/- overnight at 4°C to prevent non-specific binding of antibodies.

### LNP-mediated gene targeting

#### GFP mRNA production

GFP mRNA was synthesized by *in vitro* transcription (IVT). Briefly, linearized plasmid templates with an encoded 101-nucleotide poly(A) tail were subject to IVT using MEGAscript<SUP>TM </SUP>T7 transcription kits (ThermoFisher Scientific). mRNAs were fully substituted with N1-methylpseudouridine and co-transcriptionally capped with CleanCap Reagent AG (TriLink Biotechnologies). Cellulose chromatography was used to remove dsRNA contaminants, and mRNAs were confirmed to be the expected size by gel electrophoresis

#### Lipid nanoparticle formulation

Lipid nanoparticles (LNPS) were prepared using NanoAssemblr™ Ignite™ nanoparticle formulation systems (Cytiva) by mixing lipid mixture and mRNA at a weight ratio of 40:1, flow rate ratio at 1:3 and total flow rate of 6 ml/min. The lipid mixture contains 4A3-sc8 (Echelon Biosciences), cholesterol, DMG-PEG2000, 18:1 (Δ9-Cis) PE (DOPE) (Avanti Research) at the molar ratio of 50:38.5:1.5:10 in ethanol. EGFP mRNA was dissolved in a 50mM citrate buffer at pH 4. After formulation, LNPs were dialyzed against 1X PBS for 1 h in dialysis membrane tubing with a 6-8 kDa molecular weight cutoff (ThermoFisher Scientific). LNPs were then sterilized with a 0.22 μm filter.

#### Antibody modification

Azide-free Human PDGFRβ antibody (R&D) or Rat IgG isotype control (ThermoFisher Scientific) were functionalized with dibenzocyclooctyne (DBCO) via incubation with a 5-fold molar excess of DBCO-PEG(4)-NHS ester (Vetor Lab) in anhydrous DMSO for 1 hour at room temperature. Unreacted DBCO-PEG(4)-NHS ester was using Zeba Spin Desalting Column (ThermoFisher Scientific). Antibodies were sterilized with a 0.22 μm filter. The concentration of protein was measured using a NanoDrop spectrophotometer (ThermoFisher Scientific).

#### LNP-antibody conjugation

Azide-functionalized LNPs were formulated using the NanoAssemblr™ Ignite™ nanoparticle formulation system (Precision Nanosystems). An organic phase containing a mixture of lipids dissolved in ethanol at a molar ratio of 4A3-sc8:cholesterol:DOPE:DMG-PEG:DSPE-PEG2k-Azide (50:38.5:10:0.75:0.75) was mixed with an aqueous phase (50 mM citrate buffer, pH 4) containing eGFP mRNA or anti-*YAP1* siRNA at a total lipid:RNA weight ratio of 40:1 and a flow rate ratio of 1:3. The resulting LNPs were dialyzed against 1× PBS for 2 hr. The concentration of LNPs was determined by nanoparticle tracking analysis (NanoSight, Malvern).

Following formulation, antibodies were conjugated to LNPs based on the concentration of LNPs and proteins, aiming for approximately 100 antibodies per LNP. The conjugation reaction was carried out overnight at 4°C. Conjugation efficiency was determined by size-exclusion chromatography, as previously described (PMID: 39103056). For both MFB specific antibody and IgG LNPs, the conjugation efficiencies were 91.5% and 94.3%, respectively. Conjugated LNPs were stored at 4°C for later use.

Prior to use, the anti-*YAP1* siRNA LNP was sterilized by filtration through a 0.45 μm filter. The final sterilized MFB specific antibody LNP, conjugated with anti-*YAP1* siRNA, was then added at 5 μg/ml to each well of interest for 3 days prior to formalin treatment.

### Antibody Staining + Immunofluorescence

After overnight blocking with 0.1% BSA in 1x PBS -/- at 4°C, the 0.1% BSA in 1x PBS was removed from the fixed 96-well plate of cells. This was followed by washing twice with either PBS -/- or PBS +/+ in preparation for primary antibody staining. Each primary antibody was mixed with 0.1% BSA in 1x PBS -/- to a predetermined dilution factor specific to the antibody being used. Most, but not all, primary antibodies were used at a concentration of 1:100. Diluted antibodies were added to the desired wells of the 96-well plate and were then incubated either overnight at 4°C, or for one hour at room temperature (approximately 20°C) shielded from light.

Following the initial primary antibody incubation period, the 96-well plate was washed twice with 1x PBS +/+ to prepare for secondary antibody staining. Each secondary antibody was mixed with 0.1% BSA to a dilution factor of 1:500 and Hoechst dye (Sigma-Aldrich, #94403, final concentration 1 μg/ml). Diluted antibodies were added to the desired wells of the 96-well plate and were incubated for one hour at room temperature shielded from light. The 96-well plate was then washed twice more with 1x PBS +/+. Finally, the wells were filled with either 1x PBS +/+ and stored at 4°C until needed for microscopy.

### Single Cell Preparation and scRNA Sequencing

Triple co-cultures were grown for 7 days in fibrosis medium. On day four, 0.4 mM palmitic acid and 0.4 mM oleic acid conjugated to BSA as described (3) and were then added to the fibrosis medium. On day seven, cells were subjected to PBS -/- for 20 minutes at RT. TrypLE was then added for another approximately 10 minutes at RT and the trypsin digestion was stopped with 2% serum in RPMI. Cells were suspended and pipetted several times, spun down, and resuspended in fibrosis medium. Cells were then immediately transferred to the Children’s Hospital of Philadelphia (CHOP) Single-Cell Technology core for preparation of single-cell RNA libraries.

Single-cell RNA sequencing was performed using the 10x Genomics Chromium GEM-X Single Cell 3’ v4 Gene Expression assay following the manufacturer’s protocol (10x Genomics, Pleasanton, CA). Briefly, single-cell suspensions-derived from organoids were evaluated for viability and concentration prior to loading onto the Chromium X instrument for Gel Beads-in-Emulsion (GEM) generation, enabling the barcoding of individual cells and their transcripts. Reverse transcription and cDNA amplification were carried out within the GEMs, followed by library construction, using the Chromium GEM-X Single Cell 3’ Kit (PN-1000691), Chromium GEM-X Single Cell 3’ Chip Kit (PN-1000690), and the Dual Index Kit TT Set A (PN-1000215). Final libraries were quantified, pooled, and sequenced on an Illumina NovaSeq 6000 system using the SP Reagent Kit v1.5 (100 cycles) to a depth of approximately 14,000 – 19,000 reads per cell.

### Analysis of scRNA seq data

Demultiplexed reads were processed using CellRanger v9.0.0 from 10x Genomics. Reads were aligned to the human reference genome GRCh38. Downstream processing and analysis was performed using R v4.4.0. Unique molecular identifier (UMI) count matrices were imported into Seurat (##) v5.2.0. Ambient RNA expression was corrected using SoupX (32) v1.6.2. Briefly, counts were normalized using sctransform, and cells were filtered on a library-specific basis of number of features, total counts, mtRNA content, and rRNA content. Putative doublets were identified using DoubletFinder (33) v2.0.4. Processed count data were merged, and cell clusters were assigned using the Louvain method. Additional filtering was applied to remove likely cell debris based on feature count. The final data set consisted of 35,138 cells. Using a human cell atlas of healthy and cirrhotic liver (7), initial cell type labels were assigned using Seurat’s native reference mapping method. Labels were further refined upon manual investigation of each cluster.

### Quantification and statistical analysis of microscope images

All quantitations of organoid, HSC, and MP primary cells in mono, double, or triple co-cultures are based on the averages of 8 randomly chosen images for each well of a 96 well plate, unless otherwise noted in the figure caption. All fluorescence images were taken on a Nikon Eclipse TE2000-U Inverted Fluorescence Microscope with NIS-Elements AR 3.1 software. Image acquisition parameters were kept consistent within the same experiment and all image processing was performed equally across images of the same experiment. Images were processed with Adobe Photoshop CC 2025, and figures were assembled using Adobe Illustrator CC 2025. Statistical analyses were performed with Microsoft Excel and GraphPad Prism. P values for all figures were calculated with unpaired two-tailed t tests with error bars represented by ± SD.

## Acknowledgments

We thank the members of the Muzykantov, Weissman, and Rader labs for helpful discussions. We are grateful to the Single Cell Technology Core and the High Throughput Screening Core of the Children’s Hospital of Philadelphia Research Institute for assistance in acquiring our single cell data. We thank Teodora Orendovici, PhD, the High Throughput Screening Core technical director, for helpful discussions. We thank the Penn Immunology Core for preparation of human peripheral monocytes. This work was supported by NIH grant 1R21AI183111-01(T.R) and the Arno A. Roscher Foundation (T.R.), as well as the American Society of Gene and Cell Therapy/Cystic Fibrosis Foundation Career Development Award (J.N.)

## Author Contributions

Conceptualization and experimental design, T.R., A.L., A.d.t.H.

Writing, Original Draft, T.R., A.L., A.d.t.H., K.B.

Visualization, A.L., A.d.t.H., T.R., D.S.

Writing, Review and Editing, T.R., A.L., A.d.t.H., J.N., J.M., V.M., D.W., D.S, D.J.R.

Supervision, T.R.

Single cell RNA sequencing and analysis, D.S. T.R. A.L., M.Z.

Immunofluorescence and microscopy analysis, T.R., A.d.t.H., A.L.

Tissue harvesting and biobanking, T.R.

Tissue Culturing, T.R., A.L., A.d.t.H., K.B.

LNP Antibody Conjugation, J.N., V.M.

GFP mRNA Generation and LNP Formulation, J.N., J.M., M.K., J.M.-R., D.W.

Funding acquisition, T.R., J.N.

---

**Figure S1:**
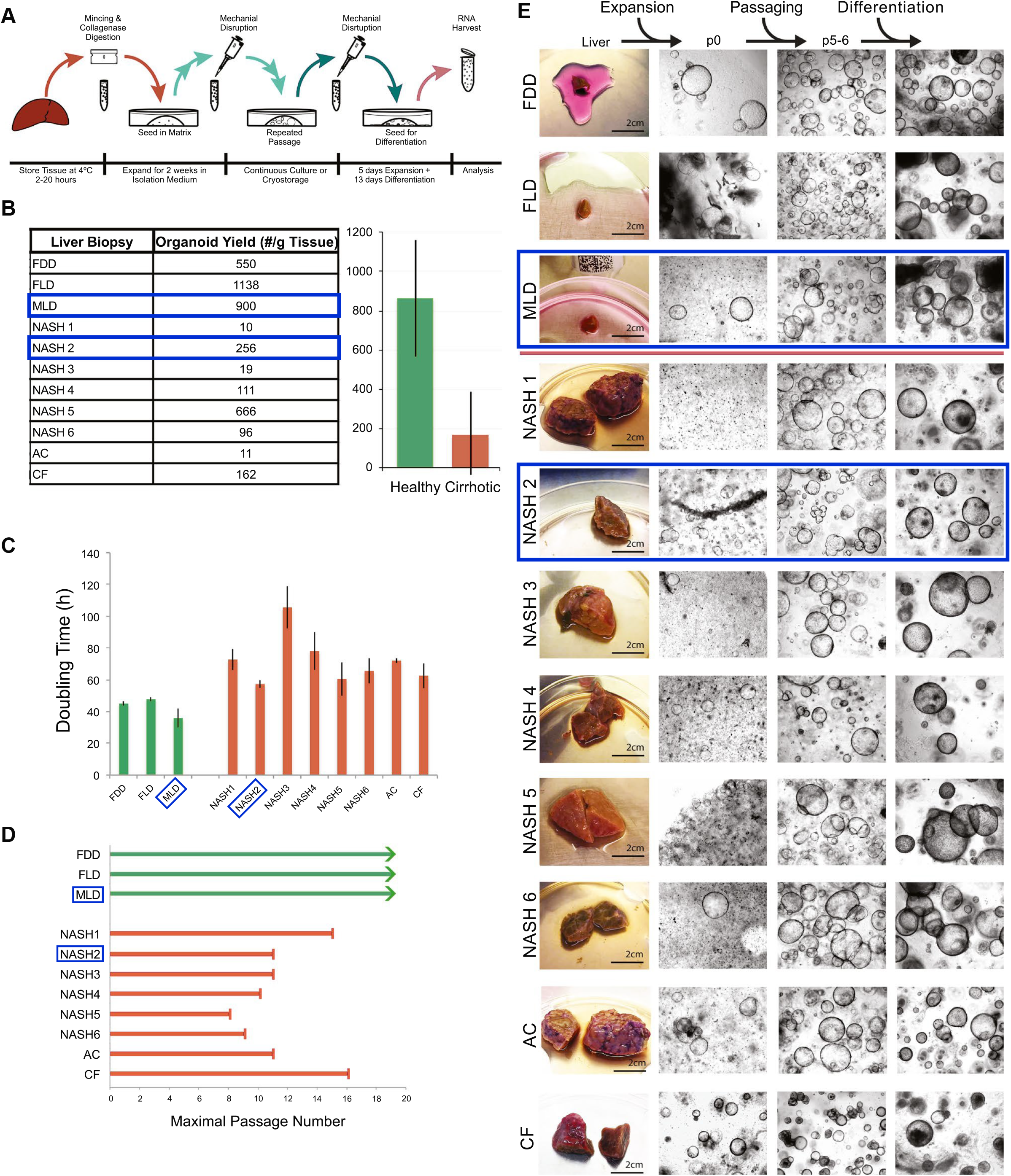
Selection of human liver-derived organoid lines for this study. Organoid lines were selected from a previously described panel of human donor liver and MASH explant liver derived organoids (3). Selected lines are boxed in blue. NASH2 is the third patient of a series of explants and denoted here as PAT3.

**Figure S2:**
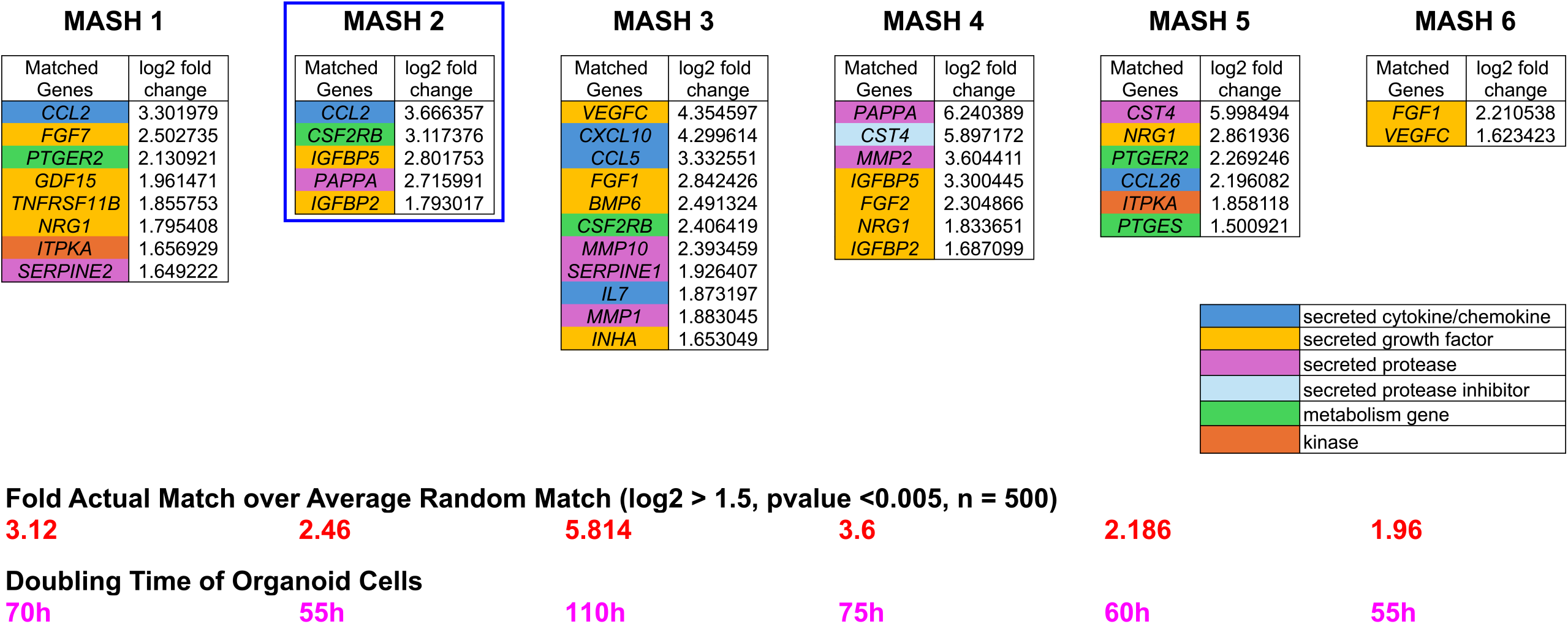
RNAseq analysis of functionally annotated senescence markers overexpressed in NASH organoids from Figure S1. Note that MASH is identical to NASH and the numbering is maintained (i.e. NASH2 = MASH2) in Figure S1 due to recent MSLD nomenclature changes. Therefore PAT3 is identical to MASH2. This line was selected for this study due to its intermediate overexpression of senescence associated factors, most of which are SASP factors. This figure shows overexpressed senescence associated genes in patient organoids. All genes shown are contained within the Sen Mayo functionally annotated gene list of 125 genes. We show fold actual match over average random match for each MASH line (n = 500, p-value<0.005), indicating true overexpression of senescent genes. Note that the slowly replicating organoid lines tend to overexpress the highest number of genes and the highest log2fold changes.

**Figure S3:**
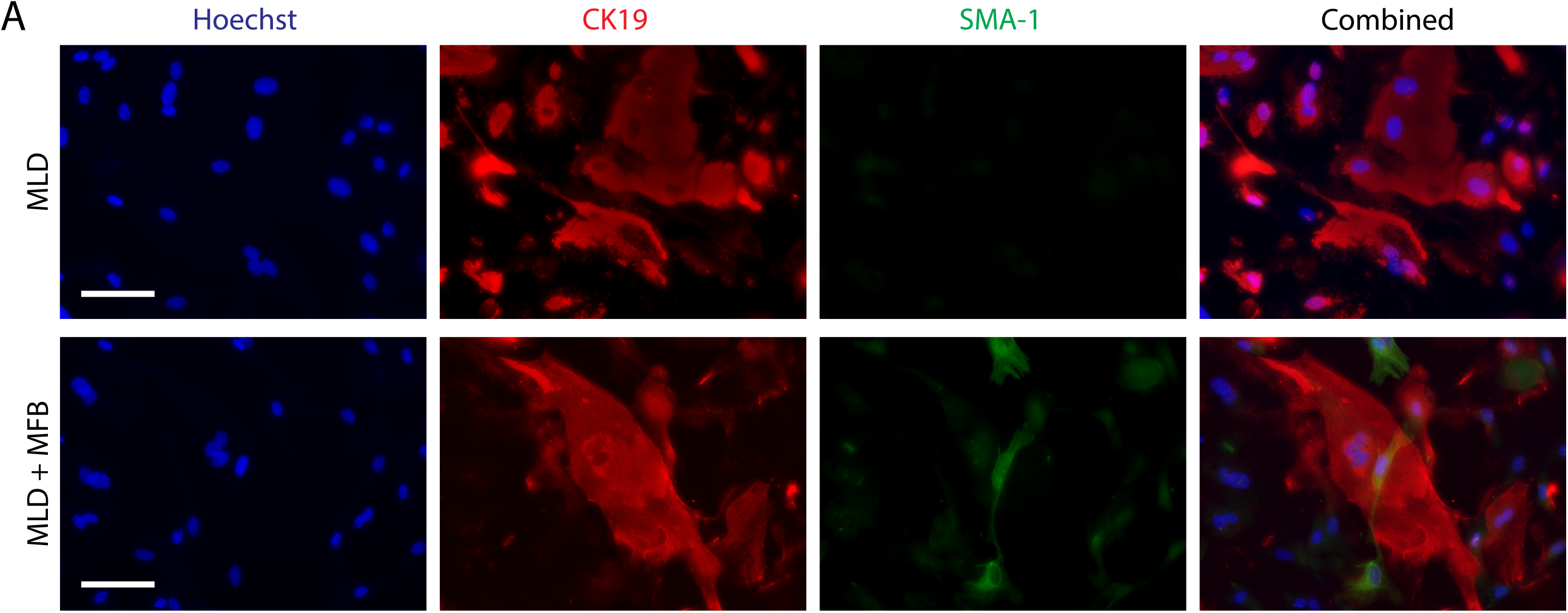
MLDs express classical cholangiocyte markers and inhibit MFB proliferation. Upper panel: MLDs express cytokeratin-19 (CK19). Lower panel: In the presence of MLD organoids, MFBs express low levels of a-smooth muscle actin (SMA-1). Notably, significantly fewer MFBs proliferate in the presence of MLD compared to PAT3 (compare lower panel to Figure 2A, lower panel, Figure 3). Scale bars, 100 µm.

**Figure S4:**
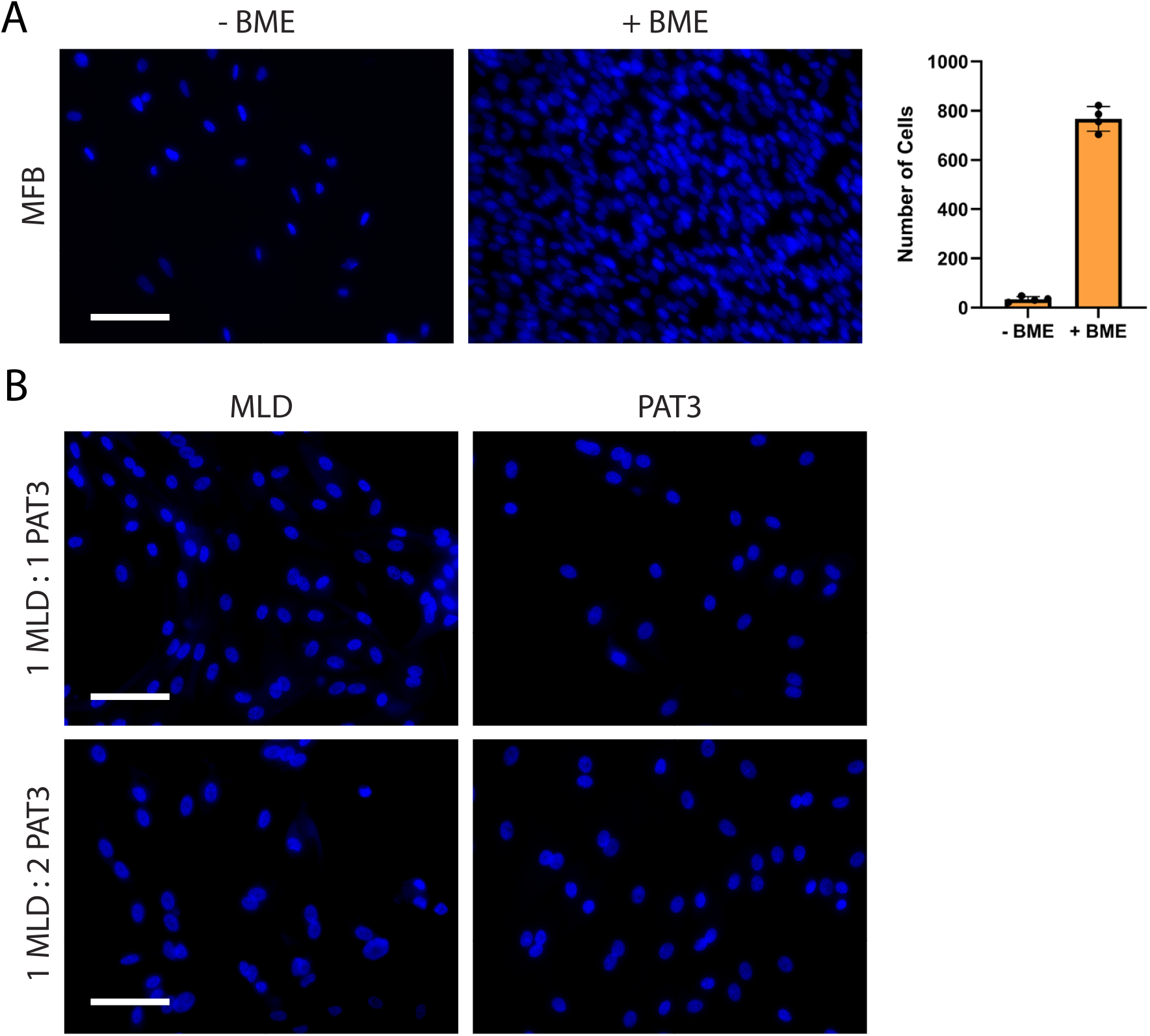
Culture conditions used to ensure equal numbers of PAT3 and MLDs at the time of MC and MFB addition, and to support formation of dense, spindle-shaped MFB aggregates. Related to figures 2 and 3. A) Strong MFB proliferation in monoculture resulting in densely packed fibrosis-like structures is fully dependent on BME (Matrigel) presence on the bottom of the culture well. Quantitative measurement of MFB replication is shown on the right (mean± SD, n = 4). Scale bars, 100 µm. B) Unlike PAT3s, MLDs do not fully arrest proliferation when exposed to differentiation media over 4-6 days, a plating ratio of 1:2 between MLD and PAT3 is necessary to ensure an equal number of cells upon full differentiation, just before the addition of MCs and MFBs. Scale bars, 100 µm.

**Figure S5:**
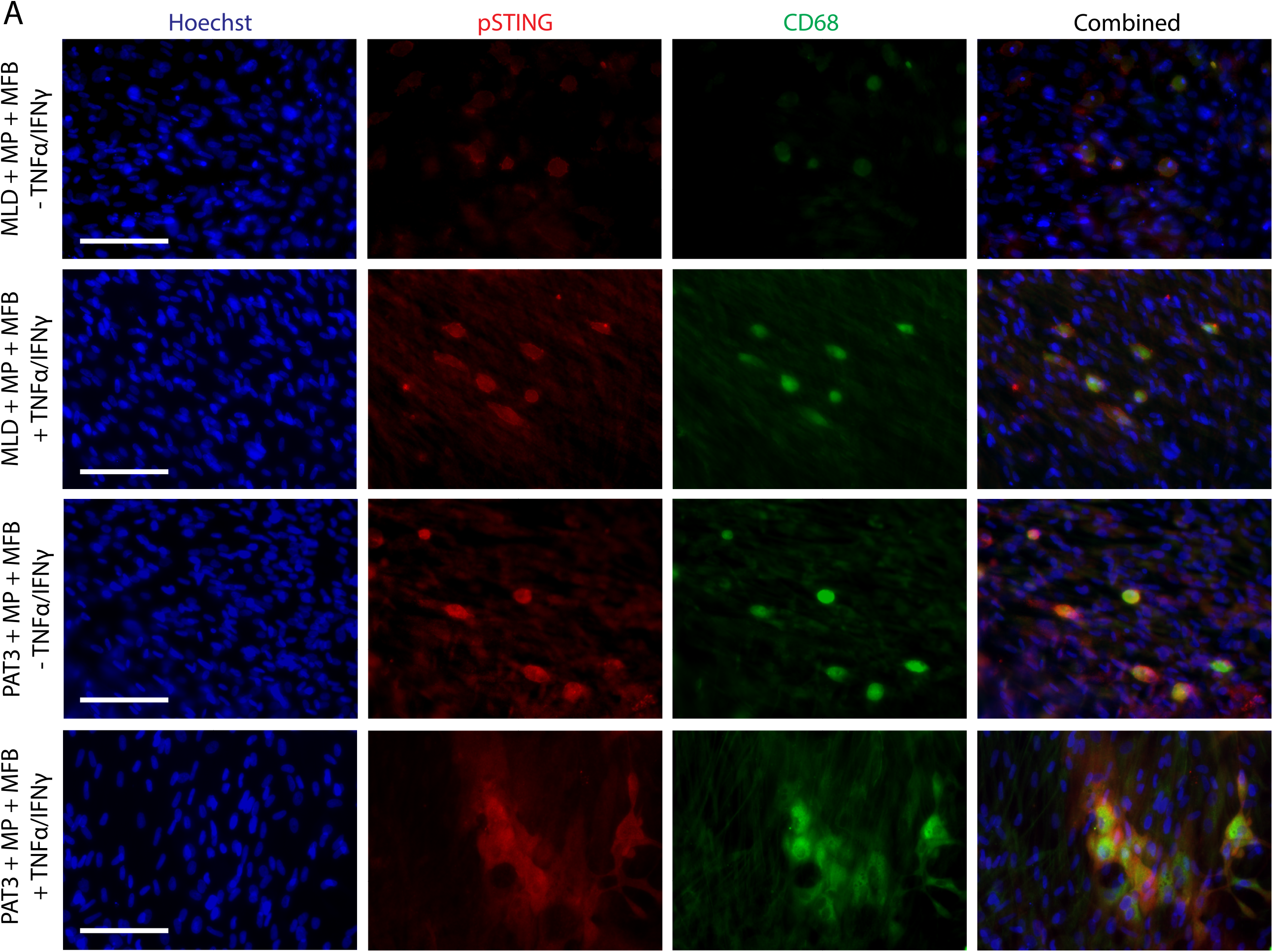
Separate pSTING and CD68 antibody stains that are only shown as merged images of branched MPs in Figure 5D. A) Upper two panels: MCs in triple co-cultures containing **MLD** do not strongly express pSTING and CD68, but the addition of inflammatory factors TNFa and IFNy strongly induces both. However, the activated MPs do not branch out. Scale bar, 100 µm. B) Lower two panels:. Unlike **MLD** triple co-cultures, PAT3 triple co-cultures spontaneously induce pSTING and CD68 expression in MPs. Additionally, TNFa and IFNy strongly induce flattening and branching of the MPs.

**Figure S6:**
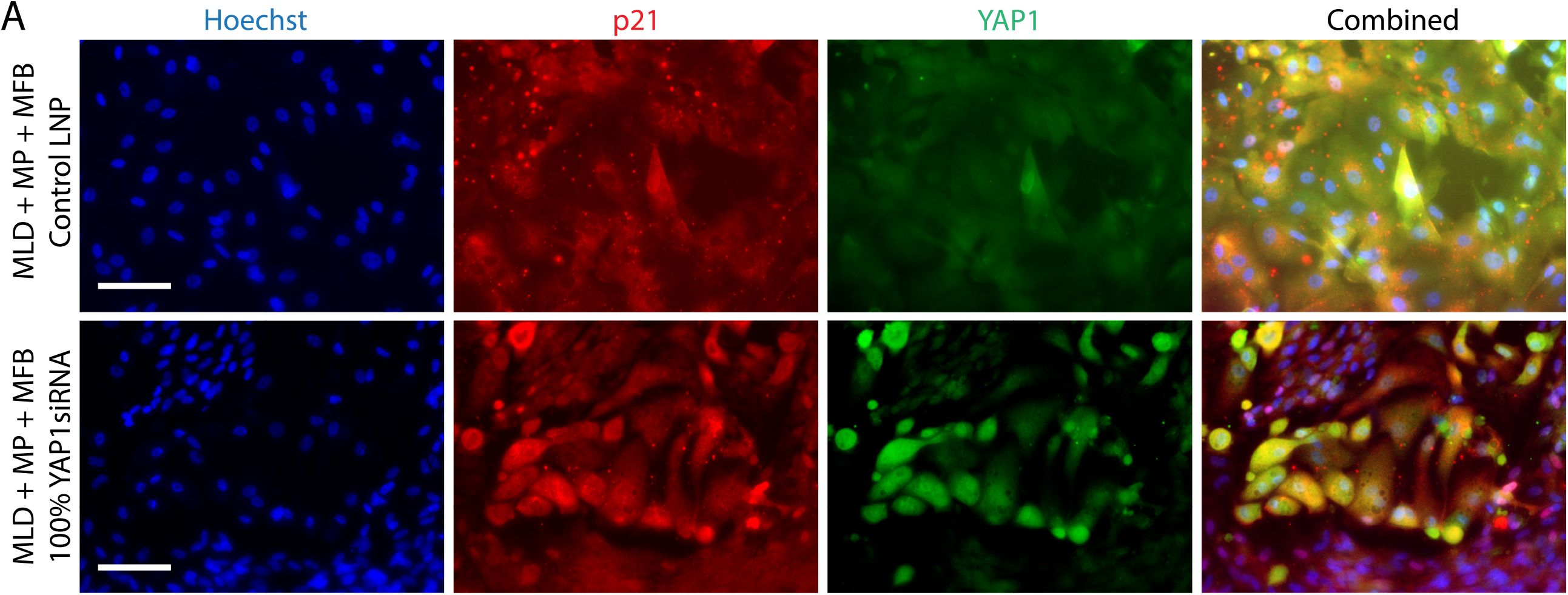
MFBs in MLD triple co-cultures exhibit similar *YAP1* knockdown levels compared to MASH liver organoid triple co-cultures. Related to Figure 7. Compared to control LNP (upper panel), YAP1siRNA LNP (lower panel) added to MLD triple co cultures induce MFB specific YAP1 protein knockdown and p21 overexpression, similar to MFBs in PAT3 triple co-culture. No change in either YAP1 or p21 is seen in organoid cells. Scale bars, 100 µm.

**Figure S7:**
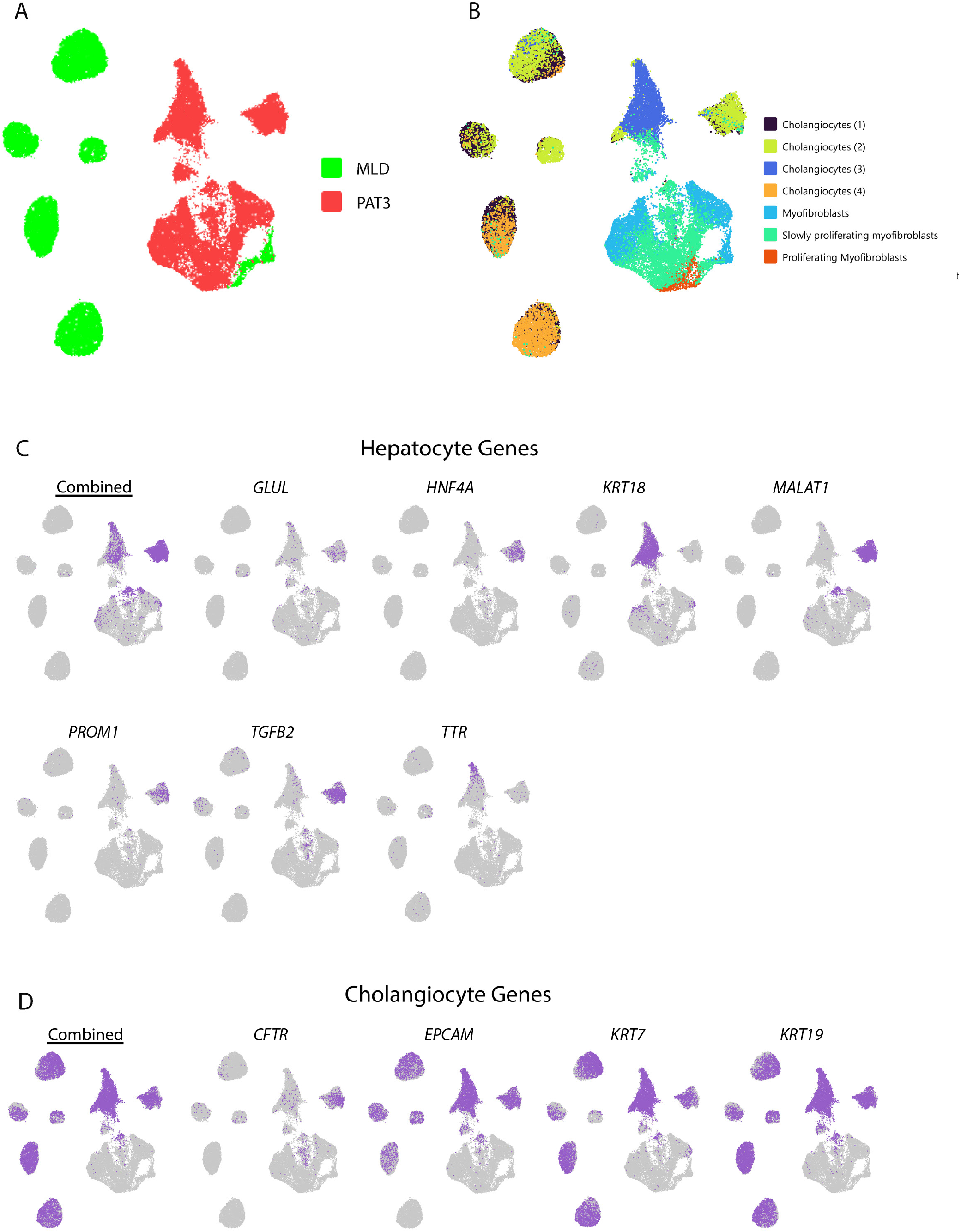
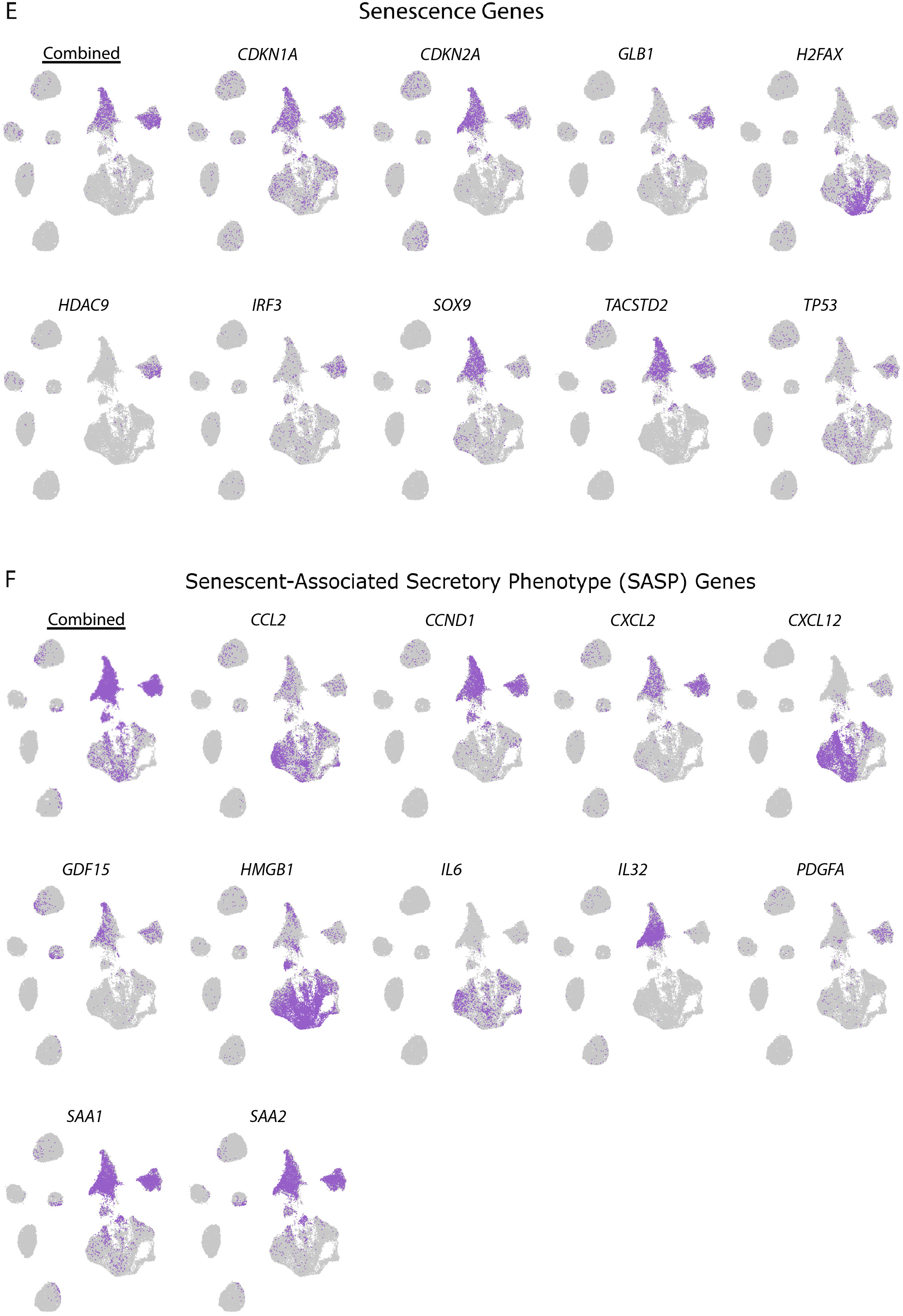
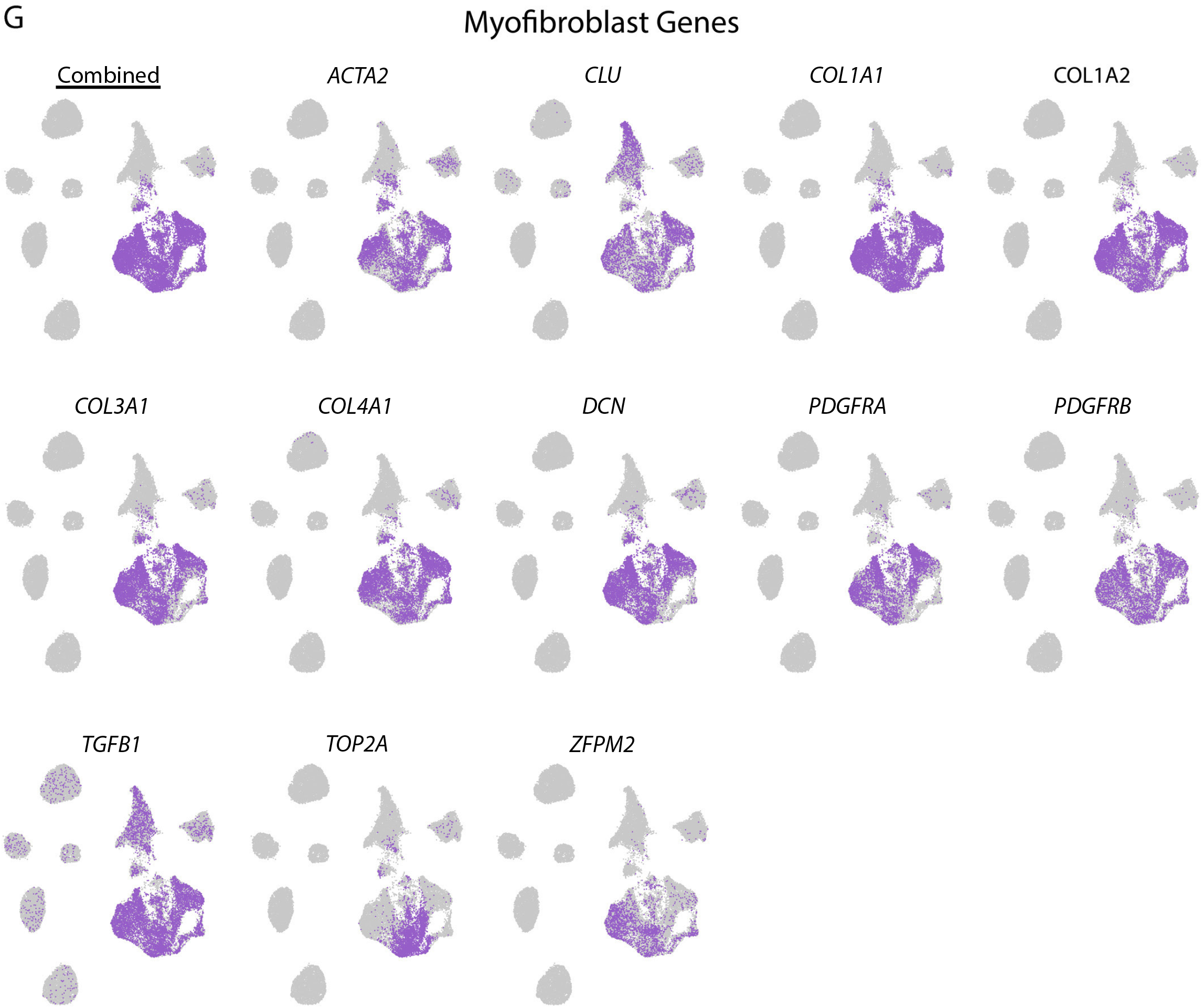
Additional single cell sequencing data. Note that combined UMAPs (underlined) of genes belonging to the same type (C, D, E, F, G) are shown as the first UMAP for each set of genes^*^. A) Combined UMAP of MLD and PAT3 triple co-cultures showing parenchymal cells and MFBs. Note that there is no meaningful overlap of any MLD and PAT3 populations. B) Cell types were predicted by unsupervised alignment of the RAW data with published single-cell data from five pooled healthy donor livers and five pooled end stage fibrotic patient livers (10). Both biphenotypic hepatocyte/ cholangiocyte populations and MFB populations are distinct between MLD control co-cultures and PAT3 fibrosis co cultures. This distinction persists for MFBs even though they originate from the same batch, and the resulting MLD and PAT3 co-cultures were grown in parallel. This indicates that PAT3s, but not MLDs, strongly stimulate MFB proliferation and activation, as functionally confirmed in Figure 3E. C) Hepatocyte genes are mostly expressed in PAT3 parenchymal cells. Some of these genes are less strongly expressed than in MLD hepatocytes, indicating a biphenotypic state. D) Cholangiocyte genes are expressed in most parenchymal cells of both the MLD and PAT3 co-cultures, However, strongly biphenotypic cholangiocyte/hepatocyte cells appear mostly in PAT3 co-cultures (see also C, above). E) Expression of genes functionally involved in senescence are predominantly over expressed in the biphenotypic cells of the PAT3 co-cultures and not in the MLD co-cultures. F) SASP genes are expressed by the senescence biphenotypic cells from C and D. Note that some MFB cells also express some SASP genes. G) MFB genes are strongly overexpressed in most subtypes of MFBs. Notably, MFBs from the MLD co-culture express some of these genes *-ACTA2* (SMA-1), *COL3A1, CLU, DCN, PDGFRA*, and *PDGFRB* - but at lower levels (see also A). Additionally, *TOP2A*, which mediates cell replication, is expressed only in a subset of MFBs. ^*^Note that MPs are missing as they were not compatible with the single cell procedure.

